# Copy number variation facilitates rapid toggling between ecological strategies

**DOI:** 10.1101/2025.07.22.666191

**Authors:** Farah Abdul-Rahman, David Gresham

## Abstract

A central question in evolutionary biology is how different types of mutations shape ecological strategy. Whereas both single nucleotide variants (SNVs) and copy number variants (CNVs) can alter gene dosage, it remains unclear whether CNVs confer unique adaptive advantages. Here, we show that CNVs do not mediate adaptive tracking in fluctuating environments, but instead act as reversible toggles between ecological strategies. Using a dual-fluorescent CNV reporter system in *Saccharomyces cerevisiae*, we tracked CNV dynamics at two transporter loci during long-term evolution under static and fluctuating nitrogen limitation. CNVs arose with high repeatability in static conditions but showed dampened or divergent dynamics in fluctuating environments, suggesting they do not track environmental change. Instead, we found that the ratio of copy number between the two loci, but not copy number at either locus alone, predicted ecological strategy: imbalanced CNV ratios defined specialists, while balanced ratios defined generalists. Evolution in static environments favored specialists whereas fluctuating environments favored generalists. Applying this framework to over 3,000 sequenced yeast genomes, we found generalist CNV signatures in both wild and domesticated strains, but specialist signatures exclusively in domesticated strains. These findings introduce a generalizable framework for predicting ecological strategy from genome structure, positioning CNV ratios as a molecular signature of niche breadth across evolutionary and ecological contexts.

## Introduction

A central goal in evolutionary biology is to infer how mutations influence phenotypic outcomes and fitness. However, the factors that bias evolutionary change toward particular types of mutations, when alternative mutations could produce similar phenotypes, remain poorly understood. Single nucleotide variants (SNVs) and copy number variants (CNVs) can both alter gene dosage (Kazem et al. 2025; Tomanek and Guet 2022; Spealman et al. 2025), yet the mechanisms and rates underlying their generation are distinct, necessitating the development of separate frameworks to study their evolutionary impacts (Kimura 1985; Ohno 1970). In contrast to SNVs, CNVs have high reversion rates (Jeffares et al. 2017), exclusively impact gene dosage, and typically confer large fitness effects. Whereas CNVs have been extensively studied for their influence on molecular and morphological phenotypes (Montanucci et al. 2023; Bai et al. 2016; DeBolt 2010), their role in shaping ecological strategies, such as specialization and generalization across multiple environmental axes (Xue, Sartori, and Leibler 2019), remains underexplored.

CNVs arise through gene amplifications or deletions, in which a single mutational event can result in either loss, duplication, triplication, or even greater copy number expansion, depending on the mechanism of generation (Hastings et al. 2009; Gresham et al. 2010). This can lead to rapid and substantial changes in fitness enabling organisms to adapt to a wide array of environments (Gyorfy et al. 2015; Gao et al. 2016). For example, the ribosomal DNA (rDNA) locus exhibits copy number variation across all domains of life, directly influencing organismal growth profiles. In bacteria, rDNA copy number is closely associated with maximal growth rate, serving as a predictor of whether a species adopts an oligotrophic or copiotrophic ecological strategy (Klappenbach, Dunbar, and Schmidt 2000; Roller, Stoddard, and Schmidt 2016).

Similar findings have been observed under stressful conditions and intraspecifically in eukaryotic microbes such as *Saccharomyces cerevisiae* (Thornton et al. 2024). The direct relationship between rDNA copy number and growth is intuitive, given its direct influence on protein translation capacity. However, whether CNVs at non-ribosomal loci also play a role in determining ecological strategy remains unknown, despite their occurrence across broad classes of genes.

Adaptation to nutrient-limited environments often involves gene amplification of the corresponding transporter gene, enabling increased uptake of scarce resources. These gene amplifications may promote ecological specialization, whereby cells enhance performance in a specific nutrient regime at the expense of flexibility in others. In nutrient-limited chemostats, *S. cerevisiae* repeatedly evolves amplifications of transporter genes for carbon, nitrogen, sulfur, and phosphate sources (Hong and Gresham 2014; Gresham et al. 2010; Payen et al. 2014; Gresham et al. 2008; Brown, Todd, and Rosenzweig 1998; Kao and Sherlock 2008; Hansche 1975). Similarly, in pathogenic and commensal *Candida* species, CNVs affecting genes encoding the efflux pumps CDR1/CDR2 (Todd and Selmecki 2020) and MDR1 (multidrug resistance transporter) (Costa et al. 2017) contribute to antifungal resistance, a form of specialization in response to drug exposure. This pattern of CNV-driven variation in transporter genes extends beyond fungi to bacteria and even vertebrates, such as the amplification of ABC transporters in *Streptococcus pneumoniae* (Silva and Khare 2024), amplifications in the citrate-succinate antiporter in *Escherichia coli* during long-term experimental evolutions (Blount et al. 2012) and SLC transporters in humans (Schaller and Lauschke 2019). The recurrence of this phenomenon across systems likely reflects the ecological importance of transporters as they are the primary interface between a cell and its environment and are not constrained by strict stoichiometric relationships. As a result, increasing transporter numbers can provide a rapid mechanism for improving acquisition of scarce nutrients with implications for niche specialization.

The rapid generation (Otto and Whitton 2000; J. Zhang 2003) and reversion of CNVs (Jeffares et al. 2017), coupled with their key role in adapting to diverse suboptimal environments, raises the question of whether they are uniquely capable of mediating adaptive tracking in periodically fluctuating conditions by dynamically altering a population’s genetic composition to continuously match environmental changes (Levins 2020; Xue, Sartori, and Leibler 2019). Whereas adaptive tracking has been observed over ecological timescales (Abdul-Rahman, Tranchina, and Gresham 2021; Rudman et al. 2022), directly observing it over evolutionary timescales remains formidable due to the technical challenges of achieving the high temporal resolution necessary to observe CNV allele frequency fluctuations over extended periods.

To address this challenge, we designed a dual-fluorescent CNV reporter system, using the tractable genetic model system *S. cerevisiae*, enabling us to simultaneously track the generation and selection of CNVs at two distinct loci at the single-cell level. We analyzed CNV dynamics at three nitrogen transporter genes in pairwise combinations - *GAP1*, *PUT4*, and *MEP2*, which are known to undergo gene amplification in specific nitrogen-limited conditions. We experimentally evolved the reporter strains in static and periodically fluctuating conditions providing the resolution to identify CNV-specific genotype-by-environment (G × E) and epistatic (G × G × E) interactions.

Contrary to our expectations, fluctuating conditions did not demonstrate CNV-mediated adaptive tracking. Instead, we found that the ratio of copy number at two transporter loci, representing distinct nitrogen environments, provided a genomic predictor of ecological strategy: imbalanced CNV ratios defined specialists, whereas balanced CNV ratios defined generalists. This framework conceptualizes CNV balance as a molecular signature of niche breadth across two environmental axes. Using experimental evolution we found that static conditions favored the emergence of specialists whereas fluctuating conditions favored the emergence of generalists. Applying this framework to over 3,000 sequenced *S. cerevisiae* genomes, we found generalists in both domesticated and wild strains, whereas specialists were restricted to domesticated isolates, likely reflecting the more stable environmental regimes experienced by the organism in human-associated settings.

Our findings shed light on the unique advantages of CNVs in regulating gene dosage in transporter genes. Contrary to our initial expectation that CNVs would facilitate adaptive tracking, we propose that they instead promote rapid transitions between ecological strategies in ways that SNVs cannot, making them essential for organismal responses to broad shifts in environmental regimes. By integrating CNVs into a specialist-generalist ecological framework, our findings highlight how gene dosage plasticity contributes to ecological adaptation. Our findings provide a novel connection between genomic structure and ecological function, suggesting a predictive model for understanding microbial evolution, engineering synthetic organisms, and reconstructing extinct species’ ecological strategies.

## Results

### A dual-fluorescence CNV reporter system tracks evolutionary dynamics at two loci simultaneously

We sought to quantify the dynamics of *de novo* CNVs at two loci simultaneously in different selective conditions. However, tracking CNV dynamics at two loci using molecular methods presents a significant hurdle as it requires isolating and sequencing numerous individual clones over multiple successive time points. To overcome this challenge, we developed a dual-fluorescence CNV reporter system that reports on gene copy number at two transporter gene loci simultaneously by expanding on methods previously established for individual loci (Lauer et al. 2018; Zhou et al. 2024; Chuong et al. 2025). We inserted two distinct fluorescent genes, encoding mCitrine and mCherry, proximate to two transporter genes known to undergo repeated amplification and selection in static nitrogen-limited environments (Hong and Gresham 2014). We constructed two dual-fluorescence reporter strains; in both strains, the mCitrine gene was inserted next to *GAP1*, which transports the nitrogen source glutamine, whereas the mCherry gene was inserted adjacent to either *MEP2*, which transports the nitrogen source ammonium sulfate, or *PUT4*, which transports the nitrogen source proline (Magasanik and Kaiser 2002). We also constructed haploid control strains with either one or two copies of mCitrine or mCherry genes inserted at neutral loci, which are not expected to evolve CNVs. Using these control strains, we confirmed that the gene copy number can be distinguished by quantitative fluorescence analysis for both fluorophores (**Supplementary figure 1A**).

We performed experimental evolution in chemostats in two types of conditions: static conditions with a single type of nitrogen-limitation, and fluctuating conditions in which the media alternated between two types of nitrogen-limitation every eight generations. The fluctuation period was chosen as CNVs form during mitotic division, requiring fluctuation periods larger than the generation time for directional selection to act on CNVs. For the *GAP1/PUT4* dual-fluorescence CNV-reporter, we used static conditions with either glutamine-limitation or proline-limitation. For the *GAP1/MEP2* dual-fluorescence CNV-reporter, the static conditions were either glutamine-limitation or ammonium sulfate-limitation. The fluctuating environments for each of the reporter strains alternated between the two respective static conditions. In all conditions, the media contained an equal molarity of nitrogen ensuring that it was only the molecular form of nitrogen that differed. All populations were evolved for a total of 260 generations with an effective population size of *N_e_ ≈* 3 × 10^7^ (**Figure 1A**). As expected, control strains did not show appreciable changes in fluorescence in either static or fluctuating conditions (**Supplementary figure 1B**).

**Figure 1.**
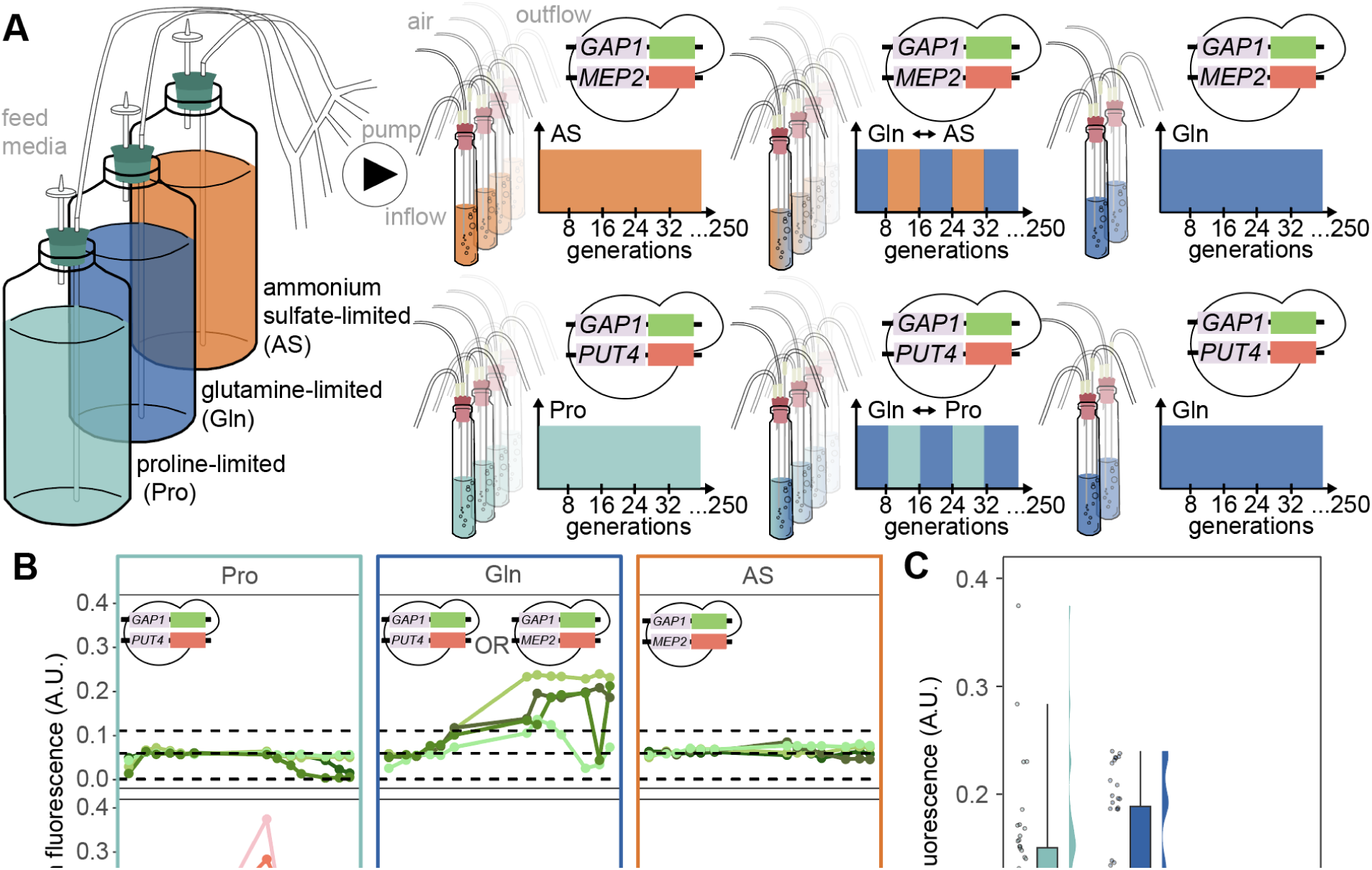
A model system for studying adaptive evolution of CNVs at two transporter loci simultaneously. (**A**) Schematic showing experimental design for both static and fluctuating conditions in continuous culture and the respective genotypes of the dual-fluorescent CNV reporter strains. (**B**) Population-level CNV dynamics are tracked using mean population fluorescence of mCitrine (top) and mCherry (bottom) in static nitrogen-limitation with the nitrogen source being either proline, glutamine or ammonium sulfate. The CNV reporter genotype in each condition is shown in each panel and the horizontal dashed black lines represent the mean fluorescence for the mCitrine and mCherry controls for 2-copy, 1-copy and 0-copies (**Supplementary Figure 1B**) respectively from top to bottom. (**C**) A summary of the overall amplification potential of each transporter is visualized using raincloud plots of aggregate mean fluorescence values from panel **B**, reporting on *PUT4*, *GAP1*, and *MEP2*, under static conditions in which their amplification is expected to provide the greatest benefit: proline-limitation, glutamine-limitation, and ammonium sulfate-limitation, respectively.

### Transporter genes exhibit locus-specific CNV dynamics in static environments

The three transporter genes *GAP1*, *PUT4*, and *MEP2*, are known to undergo gene amplification in static glutamine-, proline-, and ammonium sulfate-limited chemostats, respectively. However, the dynamics of generation and selection at these three loci has not been directly compared. Using quantification of mean population fluorescence (**Methods**) we confirmed previous observations in which *GAP1* copy number increases in glutamine-limited chemostats starting around 70 generations (**Figure 1B**). Notably, this was observed using both the *GAP1*/*PUT4* and *GAP1*/*MEP2* dual-fluorescence reporter strains, indicating that introducing a second fluorescent CNV reporter does not alter *GAP1* CNV dynamics. In static proline-limited chemostats, two populations showed evidence for selection of *GAP1* deletions consistent with a trade-off between fitness in glutamine-limitation and fitness in proline-limitation. By contrast, *GAP1* does not seem to have a similar fitness trade-off in ammonium sulfate-limitation as no populations show evidence for loss of *GAP1*.

In static ammonium sulfate-limited chemostats, transient increases in mCherry fluorescence suggest that *MEP2* undergoes copy number increases that do not reach appreciable levels in population frequency. By contrast, in proline-limited chemostats, mean mCherry fluorescence rose sharply starting at generation 70, suggesting that selection of *de novo PUT4* amplifications behave similarly to *GAP1*.

Copy number increases at *GAP1* and *PUT4* likely exceed gene duplications as mean fluorescence for both reporters exceeded mean fluorescence levels of the 2-copy controls. *PUT4* copy number dynamics are unique in comparison to *GAP1* as they showed a sharp decline in copy number after generation 200 (**Figure 1B**). This may indicate that new beneficial non-CNV mutations arose in the population or that high copy number of *PUT4* depletes environmental proline, thereby altering the selective pressure.

In static conditions, transporter genes exhibited varying degrees of amplification. The *PUT4* reporter showed the highest achieved fluorescence, the *GAP1* reporter reached intermediate levels, and the *MEP2* reporter displayed the lowest fluorescence. This suggests that *PUT4* has the greatest potential for allele expansion, whereas *MEP2* exhibits the most limited amplification capacity (**Figure 1C**), highlighting that CNV generation and selection dynamics are locus-specific.

### Fluctuating environments dampen CNV dynamics compared to static conditions

To determine whether CNVs, given their rapid rates of generation and reversion, are uniquely suited to mediate adaptive tracking, we studied their dynamics under environmental fluctuations by alternating the nitrogen source in nitrogen-limited chemostats. We first assessed the mean fluorescence of each CNV reporter and found that CNV dynamics in response to environmental fluctuations are not simply an average of dynamics of the two static conditions and that dynamics are variable between populations. If CNVs mediate adaptive tracking, the expectation would be that amplification frequency would oscillate to match that of the environmental fluctuation. However, there was no evidence of this phenomenon in any of the replicates fluctuating between glutamine-limitation and proline-limitation. In the majority of populations (replicate 1, 2 and 4) mean fluorescence appears to remain under the 2-copy control threshold. In one population, it appears that the fluctuation slowed the timing with which CNVs increase in frequency (replicate 3). Similarly, fluctuating between glutamine- and ammonium sulfate-limitation results in either transient increases in amplifications (replicate 4) or no appreciable amplifications (replicates 1, 2, and 3) (**Supplementary figure 3**).

To quantify CNV dynamics at the single-cell level, we conducted a detailed analysis of single-cell fluorescence profiles (**Methods**). We focused on glutamine-limitation, proline-limitation and the fluctuation between them since *GAP1* and *PUT4* exhibited the greatest CNV dynamics (**Figure 2**), in contrast with *MEP2* which showed much fewer changes in copy number (**Supplementary figure 4 (bottom)**). In static glutamine-limitation, the initial rise in *GAP1* frequency is a result of lineages containing *GAP1* duplications, followed by a second wave of lineages with three copies or more. We also found that a subpopulation with two copies of *PUT4* arose in replicate 2 towards the end of the experiment suggesting that either *PUT4* amplifications have a previously unrecognized advantage in glutamine-limiting environments, or that *PUT4* amplifications frequently occur and may have risen in frequency as a result due to hitch-hiking.

**Figure 2.**
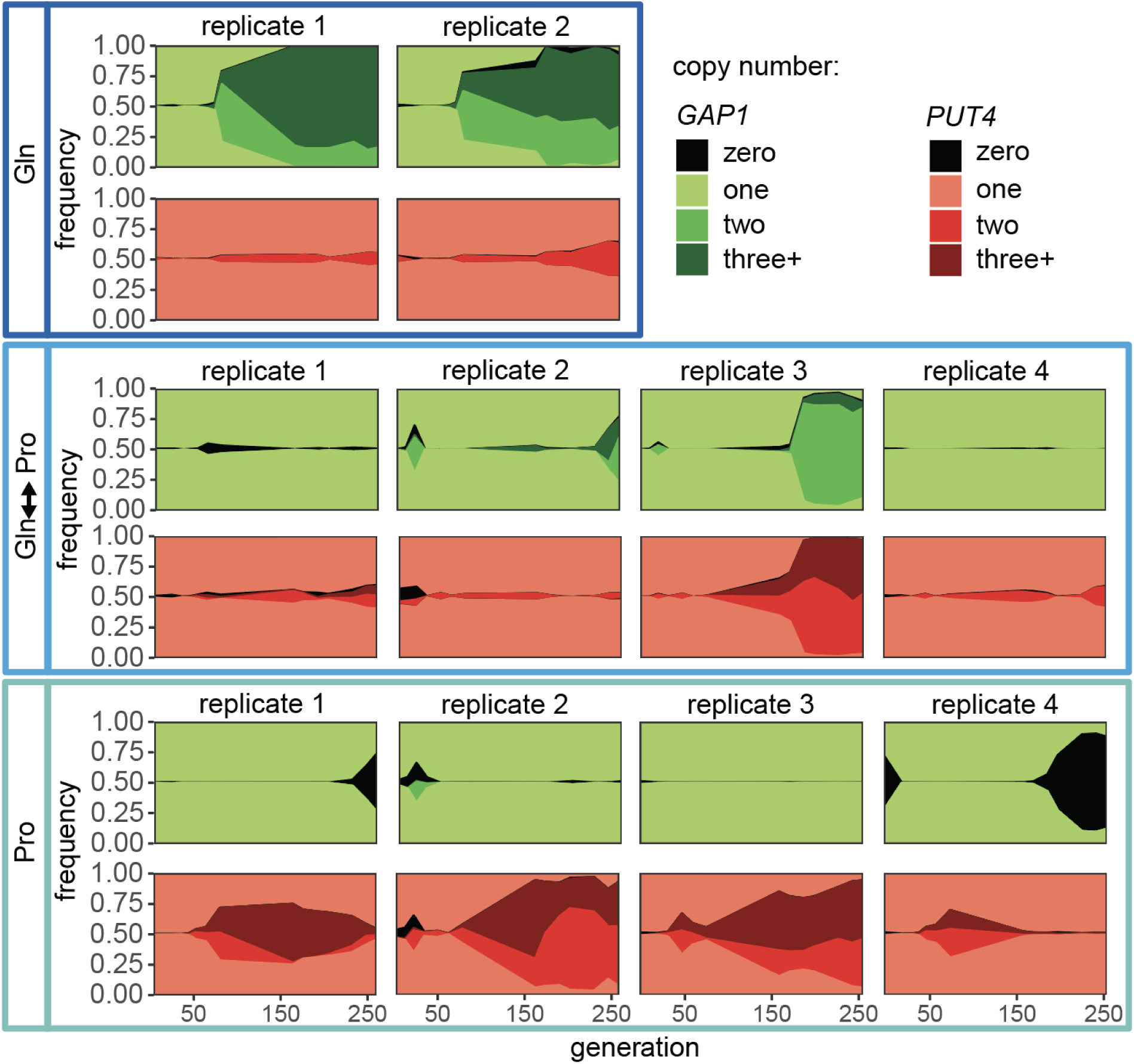
Single-cell dynamics of copy number variation at two loci in static and fluctuating environments. The proportion of the populations containing zero, one, two or three and greater copy number of either *GAP1* (green) or *PUT4* (red) is shown in static glutamine-limitation (**top**), proline-limitation (**bottom**) and the fluctuation between the two (**middle**). The frequency of each copy number class was determined by assigning either mCitrine or mCherry copy number for each cell based on gates determined using fluorescent copy number controls. Green represents *GAP1* CNVs detected using the mCitrine CNV reporter and red represents mCherry signal reporting on *PUT4* CNVs.

In the static proline-limiting condition we observed two classes of trajectories: either complete sweeps by lineages containing two or more copies of *PUT4* or the transient increase in the frequency of CNV lineages followed by return to single copy. This latter phase coincided with the emergence of a subpopulation with complete *GAP1* deletion. By contrast, fluctuating conditions appeared to suppress increases in copy number when they occurred at a single locus. This is evident in replicates 1 and 4, which consistently maintained a single copy of *GAP1* and *PUT4*, and in replicate 2, which exhibited only a transient increase in *GAP1* copy number. Interestingly, the only replicate in which copy number rose to an appreciable frequency and was nearly fixed in the population was in replicate 3. Notably, these lineages showed copy number increases of both *GAP1* and *PUT4* (**Figure 2**). Despite *MEP2* exhibiting overall dampened dynamics in static ammonium sulfate-limitation, we found a similar trend between the glutamine-to-ammonium sulfate fluctuation and the glutamine-to-proline fluctuation. Specifically, three replicates showed no changes in *GAP1* or *MEP2* copy number, whereas one replicate displayed simultaneous increases in both *GAP1* and *MEP2* loci. This further supports our observation that fluctuating conditions exhibit overall diminished CNV dynamics in comparison to static conditions (**Supplementary figure 4**), with the sole exception occurring when both loci undergo amplification simultaneously. This finding is consistent with previous reports of the gene amplification playing a more significant role in static conditions compared with a fluctuating regime that cycled between nitrogen-rich and nitrogen-poor conditions (Hays et al. 2023).

### *GAP1* and *PUT4* exhibit copy number-specific environmental and epistatic interactions

CNVs have a direct relationship between genotype and phenotype, as gene copy number directly influences the magnitude of the protein function. For instance, when *GAP1* is deleted (zero copies), cells have a significantly reduced ability to import glutamine (Grenson, Hou, and Crabeel 1970), likely resulting in a significant fitness reduction when glutamine is the sole nitrogen source. Conversely, increasing the copy number of *GAP1* is expected to improve glutamine uptake and therefore increase fitness in glutamine-limiting environments. However, this relationship can be altered by regulatory processes, such as dosage compensation or context-dependent changes in gene expression (Spealman et al. 2025). To study how copy number impacted fitness in our evolution experiments, we calculated the change in abundance of each copy number genotype between the first and final time point, a commonly used fitness estimate in a variety of barcode-based assays for short-term competition assays (Robinson et al. 2014) and long-term experimental evolution (Levy et al. 2015), for each copy number genotype combination (e.g., 1:1, 1:2, 1:3 for *GAP1*:*PUT4*).

We find that fitness increases with copy number under conditions in which increased transporter production is expected to be advantageous, particularly when the transporter’s substrate was the sole nitrogen source. Specifically, we observed a significant relationship between *GAP1* copy number and fitness in glutamine-limitation, as well as between *PUT4* copy number and fitness in proline-limitation (**Figure 3A**). We observed a similar trend in evolution experiments at the *MEP2* locus in static ammonium sulfate-limitation, although the strength of this relationship is reduced (**Supplementary figure 5A**). Surprisingly, although no significant relationship was detected for *GAP1* in the fluctuating condition, we found a significant positive relationship between *PUT4* copy number and fitness in the fluctuating environment. We also identified a significant negative relationship between *GAP1* copy number and fitness in proline-limitation, suggesting that increased *GAP1* copy numbers can have a deleterious effect in this condition (**Figure 3A**). These results demonstrate that gene copy number can have diverse effects on fitness depending on environmental context, highlighting the role of genotype-by-environment (G × E) interactions in CNV fitness effects and the resultant dynamics.

**Figure 3.**
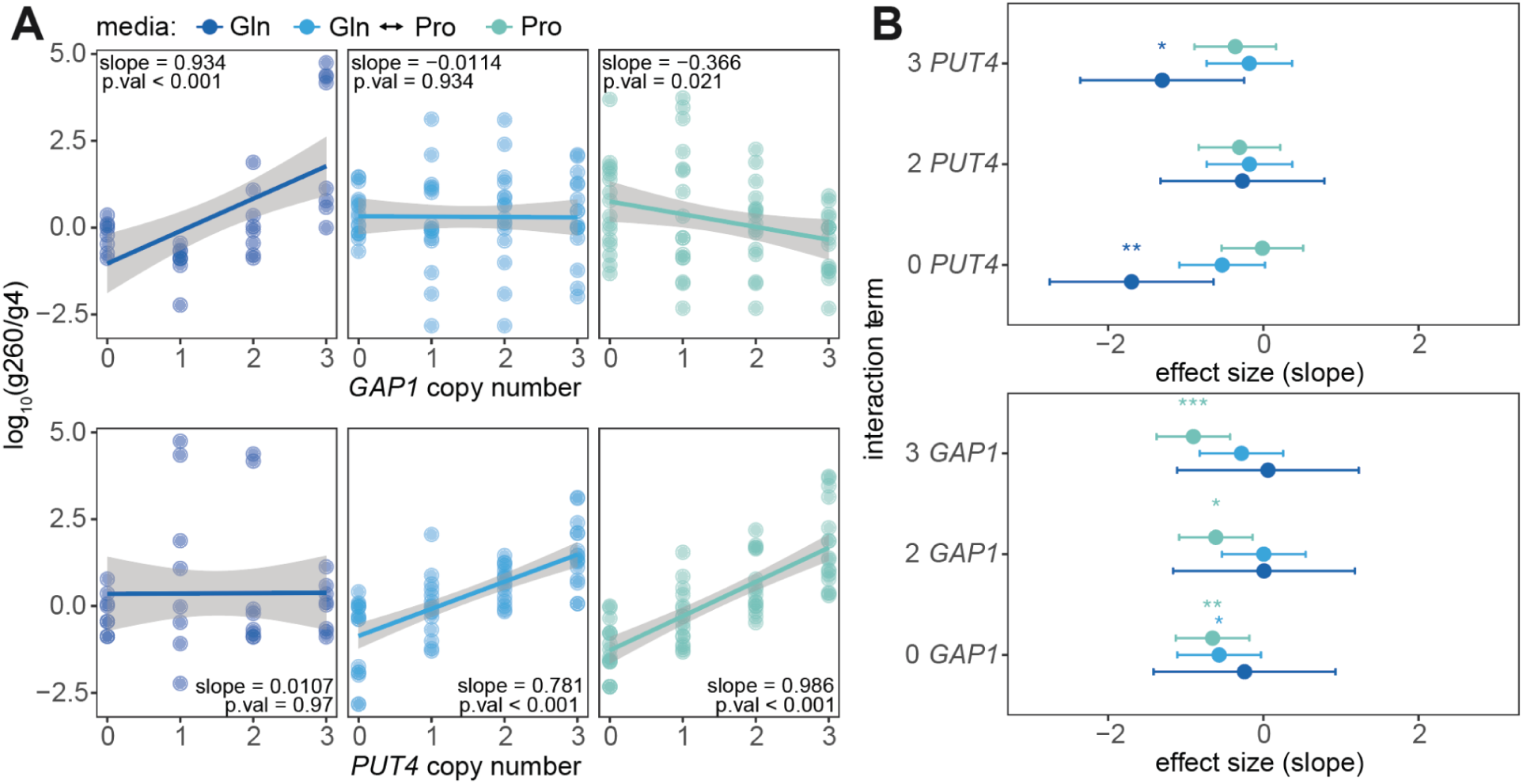
Quantification of environmental and genetic interactions during evolution experiments. (**A**) The change in ratio between generation 260 and generation 4 for each copy number variant class at the *GAP1* (top) and *PUT4* (bottom) locus in static and fluctuating environments. Increasing copy number of the relevant nutrient transporter gene results in an increase in fitness in both static environments. Increasing the copy number of a secondary transporter gene either has no effect or is slightly deleterious. In fluctuating environments, increasing the copy number of *PUT4* is beneficial whereas increasing the copy number of *GAP1* does not significantly impact fitness. (**B**) We tested whether copy number values of the secondary locus interact with the primary locus to alter fitness (G × G × E) using linear modeling. All significant interaction terms are negative and they are predominantly found in static conditions. P-values are annotated with an asterisk as follows: ‘*’ for <0.05, ‘**’ for < 0.01, ‘***’ for < 0.001.

The dual-fluorescence reporter system enables us to investigate CNV-specific epistatic interactions (G × G × E) between *GAP1* and *PUT4*. Using a linear model that includes interaction terms between the copy number of a primary locus and copy number at the secondary locus, we determined whether significant interactions occurred between the two loci in the different environments. We used a single copy of either *GAP1* or *PUT4* as the reference for our interaction terms, as our ancestral population originally contained a single copy of each. We identified a significant interaction under glutamine-limitation, in which extreme *PUT4* copy numbers (0 and 3) have a negative interaction with *GAP1* CNVs. Additionally, under proline-limitation, any variation in *GAP1* copy number negatively impacts the relationship between *PUT4* copy number and fitness. In fluctuating conditions, the only significant interaction we observed was between 0 copies of *GAP1* and variation in *PUT4* copy number. Notably, whereas G × E interactions were both positive and negative, all G × G × E (epistatic) interactions we identified were negative (**Figure 3B**). This trend was also observed for interactions between *MEP2* and *GAP1* CNVs (**Supplementary figure 5B**).

### Transporter gene copy number ratio correlates with ecological strategy

We aimed to determine whether gene copy number could predict two distinct ecological strategies: specialization and generalization. Whereas specialists and generalists are defined by their fitness profiles in different environments, identifying genetic features that reliably predict these alternate strategies has been challenging. Given the direct relationship between copy number and fitness in environments where transporters confer an advantage, CNV genotypes present a promising avenue for distinguishing ecological strategies.

Specialists exhibit asymmetric fitness profiles, achieving higher fitness in a specific nutrient environment while performing less well in one or more alternative environments. This trade-off is fundamental to specialization, as adaptation to one resource often comes at the expense of performance on others. By contrast, generalists display balanced fitness profiles, maintaining equivalent fitness across two or more environmental conditions. Generalists can follow one of two strategies: (1) no-cost generalism, in which they grow equally well in multiple environments without a fitness trade-off, or (2) costly generalism, in which they exhibit reduced efficiency across different environments (Remold 2012).

Given the direct relationship we observed between copy number at specific transporter gene loci and fitness (**Figure 3A**), gene copy number can be conceptualized as a spectrum of performance for a specific task, ranging from complete inadequacy (gene deletion) to high performance (multiple copies). Each CNV genotype occupies a position along this continuum. When considering two nutrient transporters simultaneously in a fluctuating environment a genotype’s performance extends into a two-dimensional space, representing proficiency in two distinct tasks. The position within this space defines whether an individual functions as a specialist or a generalist.

To link copy number genotype to specialization or generalization, we considered both loci simultaneously to assess fitness. Using the fitness metrics derived from evolution experiments, we compared fitness between static proline-limitation and static glutamine-limitation for each pairwise combination of copy numbers for *GAP1* and *PUT4*. As per the definition of specialism and generalism, we expected generalists to display balanced fitness between the two conditions (i.e. fall along the diagonal line) and specialists to display asymmetric fitness (i.e. fall above or below the diagonal).

We first examined the relationship between copy number of either *PUT4* (**Figure 4A**) or *GAP1* (**Figure 4B**) and fitness in either static condition and found no clear relationship predictive of ecological strategy. We reasoned that because transporter gene copy number scales with fitness in the respective substrate, then asymmetric copy number may lead to asymmetric fitness, whereas balanced copy number would lead to balanced fitness. Therefore, we examined the relationship between the ratio of copy number at *GAP1* and *PUT4* and fitness in the two static environments. Consistent with expectation, we found that all genotypes with a higher *GAP1*:*PUT4* copy number ratio exhibited higher fitness in glutamine-limiting conditions than in proline-limiting conditions. Conversely, all genotypes with a lower *GAP1:PUT4* copy number ratio had higher fitness in proline-limited conditions and negative fitness in glutamine-limited conditions. However, genotypes with equal copy numbers of *GAP1*:*PUT4* (i.e. 0:0, 1:1, 2:2, 3:3) fall on the diagonal line indicating that they have balanced fitness, consistent with generalist behavior (**Figure 4C**).

**Figure 4.**
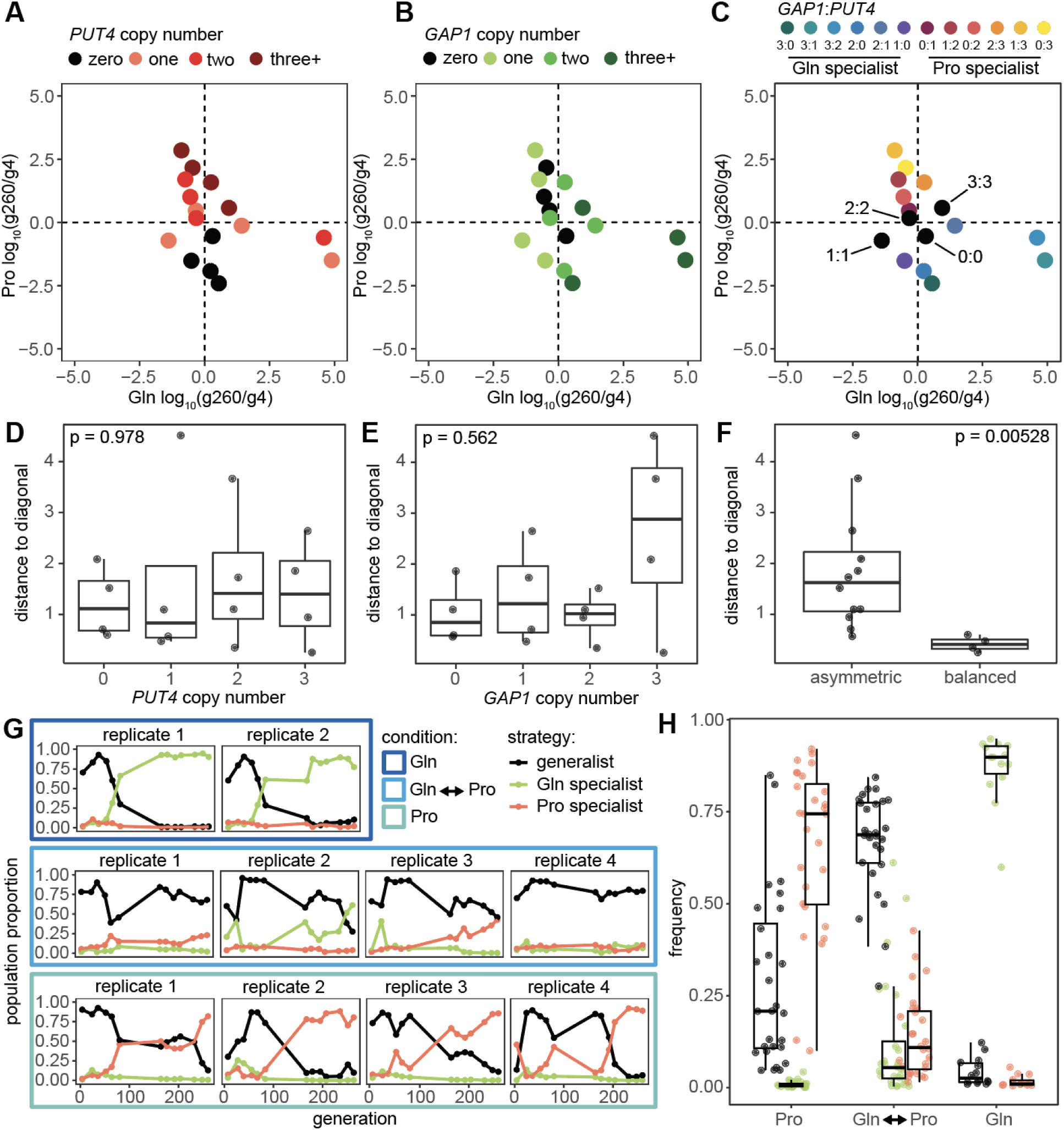
Fluctuating conditions enrich for generalists, whereas static conditions enrich for specialists. The average proportion of different CNV classes, representing every pairwise combination of *GAP1*:*PUT4* copy numbers, was compared between the two static conditions of glutamine-limitation and proline-limitation from generation 4 to generation 260. Points are colored either by (**A**) *PUT4* copy number, (**B**) *GAP1* copy number, or (**C**) the ratio between them with expected generalists colored in black. The perpendicular distance from the diagonal (x = y) was used to test whether each feature predicted ecological strategy. A Kruskal-Wallis test was used to assess significance. We compared distances based on (**D**) *PUT4* copy number, (**E**) *GAP1* copy number, and (**F**) the *PUT4*:*GAP1* copy-number ratio. Based on definitions in C, the combined proportions of specialists and generalists were mapped over time (**G**) in glutamine-limitation (top), proline-limitation (bottom), and fluctuation (middle) for replicate populations . (**H**) The frequency of specialists and generalists in each population at all generations greater than 100.

Using this representation of fitness space, we tested whether balanced CNV genotypes (in which *PUT4* and *GAP1* copy numbers are equal) lie closer to the line of equality than unbalanced genotypes. For each genotype, we computed its perpendicular distance from the diagonal for three different classes: *PUT4* copy number (**Figure 4D**), *GAP1* copy number (**Figure 4E**), and CNV ratio (**Figure 4F**). No individual *PUT4* or *GAP1* copy-number value was significantly closer to the diagonal, however, genotypes classified as having balanced copy number consistently had significantly smaller distances to the line of equality than those classified as unbalanced (Kruskal–Wallis test: χ² = 7.78, p = 0.00528) (**Figure 4F**). We tested whether this result was robust across time points through the evolution experiment by calculating fitness for all pairwise comparisons of generations relative to generation 4. The result remained statistically significant across comparisons (Kruskal–Wallis test: *χ²* = 37.509, *p* = 9.1E-10) despite earlier generations having minimal time to diversify (**Supplementary Figure 7B**). Our results demonstrate that the ratio between two transporter gene copy numbers is predictive of ecological strategy in our evolution experiments.

### Fluctuating conditions favor selection of generalists, whereas static conditions favor specialists

Using the relationship between copy number ratio and ecological strategy, we categorized cells in each evolving population as either specialists or generalists and tracked their frequency over time in each condition to determine whether specific ecological strategies are favored under different environmental regimes. We found that glutamine specialists dominated in glutamine-limiting conditions, comprising ∼90% of the population by the final time point, whereas proline specialists dominated in proline-limiting conditions, reaching ∼80% of the population across all replicate populations.

By contrast, generalists were the predominant genotype class in fluctuating environments throughout most of the time course, though exceptions were observed. In replicate 2, for instance, glutamine specialists surpassed generalists in frequency (∼55%) toward the end of the experiment. Interestingly, this population exhibited multiple transient increases in glutamine specialists, suggesting that these fluctuations may be linked to the duration of exposure to glutamine. In replicate 3, although generalists remained dominant for most of the time course, proline specialists reached an equivalent frequency at the final time point (**Figure 4G** and **Figure 4H**). This likely resulted from a subpopulation with a 2:2 *GAP1*:*PUT4* copy number ratio acquiring an additional *PUT4* copy, leading to a subpopulation with a 2:3 ratio (**Figure 2**).

Importantly, given that all populations were seeded with a generalist ancestor (1:1), we found that in static conditions there was a rapid shift from generalism to specialism (1:2, 1:3), whereas in fluctuating conditions, we found that populations either maintained the generalist genotype (1:1) or shifted to other generalist genotypes (2:2 or 3:3). These dynamics highlight the remarkable plasticity conferred by CNVs, allowing populations to rapidly switch ecological strategies in response to fluctuating environmental pressures.

### Specialization is exclusively associated with domesticated strains, whereas generalization is associated with both domesticated and wild strains

We sought to test whether a definition of specialist and generalist based on copy number ratio is applicable to genetically diverse strains by analyzing publicly available genomic data from the 3,034 *S. cerevisiae* project *(Loegler, Friedrich, and Schacherer 2024)*. We compared normalized read depth of the transporter genes *GAP1*, *PUT4*, and *MEP2* to the three housekeeping genes, *FBA1*, *RPL4A*, and *RPO21* for which strong selection is expected to prevent copy number variation. These controls define the expected variation in normalized read depth for single-copy genes. We found that most strains carried a single copy of the nitrogen transporters, though their read depth exhibited greater variability compared to housekeeping genes. Notably, nitrogen transporter genes displayed higher upper limits in sequence read depth, with *PUT4* showing the highest values, followed by *GAP1* and *MEP2* (**Figure 5A**). This trend mirrors our experimental evolution data, in which *PUT4* exhibited the highest fluorescence values, followed by *GAP1* and *MEP2* (**Figure 1C**). We also found a similar trend in other transporters for nitrogen, carbon, and phosphorus and other substrates (**Supplementary figure 8C-D**)

**Figure 5.**
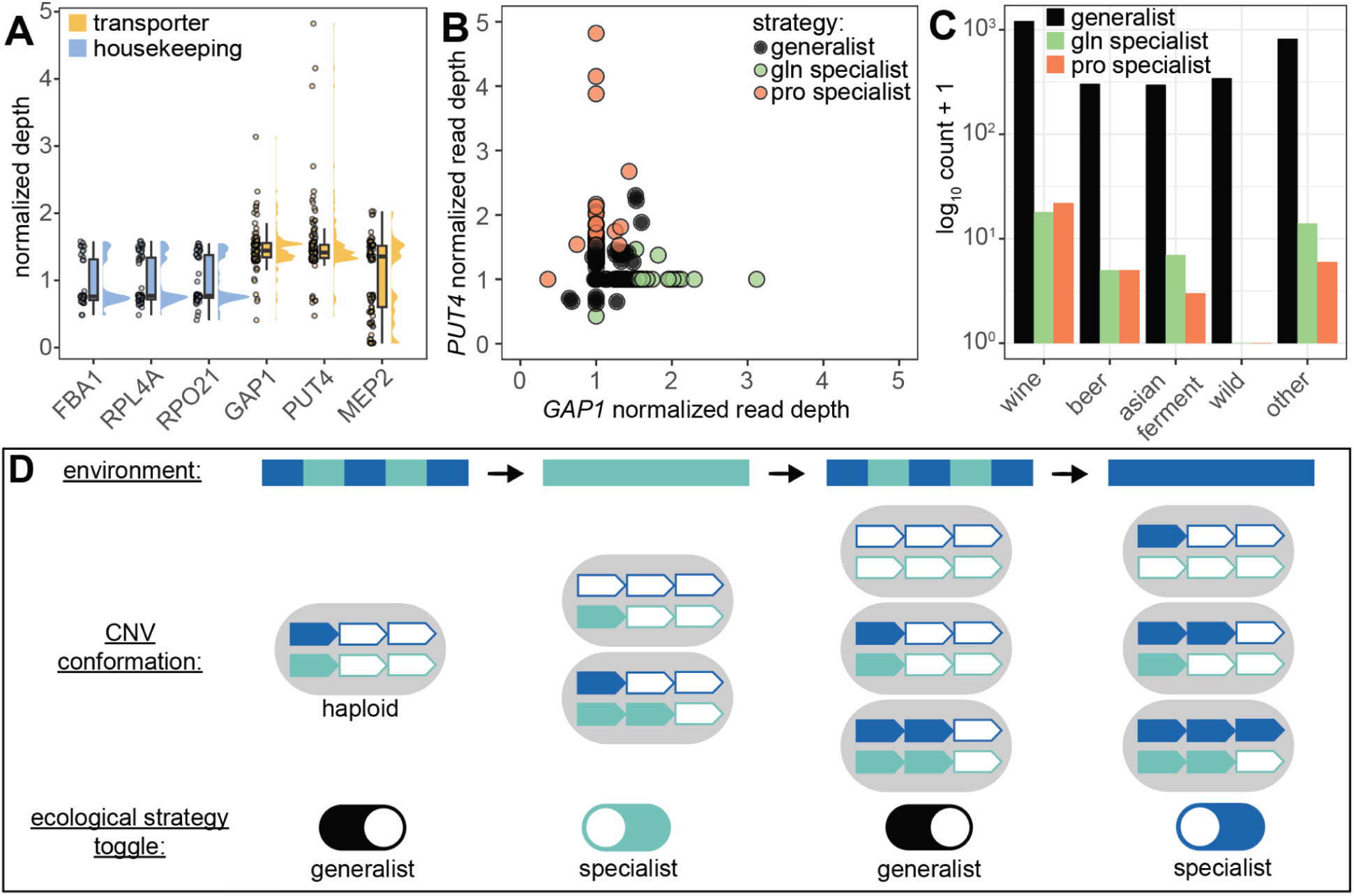
Ecological strategies predicted by transporter copy number ratio reflect strain provenance. (**A**) Normalized read depth of three housekeeping genes and three nitrogen transporter genes obtained from the 3,034 yeast strain project (Loegler, Friedrich, and Schacherer 2024) after filtering out strains with a single gene copy. (**B**) Normalized read depth for *PUT4* and *GAP1* as a proxy for gene copy number, with ecological strategy assignments determined by rounding these values and applying our experimentally derived definitions based on gene copy number ratio. (**C**) Barplots showing the number of strains falling within each ecological strategy across yeast isolate super clades. (**D**) Theoretical model of how transporter gene copy number ratio rapidly shifts cells between balanced and asymmetric fitness values to toggle between generalist and specialist strategies matching broad changes in environmental regimes.

We classified the 3,034 strains as specialists or generalists based on transporter copy number ratios, either between *GAP1* and *PUT4* or *GAP1* and *MEP2*. We found that the vast majority of strains are generalists with respect to glutamine and proline use (**Figure 5B**) or glutamine and ammonium sulfate use (**Supplementary figure 8A**). This likely reflects our observations that specialists are selected for in static conditions, whereas generalists are favored in fluctuating environments. Given that natural environments are expected to fluctuate over time, and prolonged static conditions are relatively rare, the predominance of generalists aligns with ecological expectations. We examined the provenance of generalist and specialist strains defined by copy number ratios at the three loci. We found that generalists were distributed across both domesticated and wild strain clades, whereas specialists are exclusively found among domesticated isolates derived from wine, beer, or other fermentation processes. This trend was consistent for both *GAP1:PUT4* (**Figure 5C**) and *GAP1:MEP2* (**Supplementary figure 8B**), suggesting that long-term selection in stable, resource-specific environments may have driven the evolution of specialized nutrient acquisition strategies uniquely in domesticated lineages.

Our findings using experimental evolution and analysis of natural genetic variation reveals that transporter CNVs enable individuals to toggle (i.e. switch from one discrete state to another) between generalist and specialist states through either balanced or asymmetric copy numbers at two loci. Because generalists are favored in fluctuating conditions and specialists are favored in static conditions, we propose that CNVs uniquely facilitate adaptation across broad environmental regimes, where extended periods of relative stability (i.e. static conditions) alternate with extended periods of instability (i.e. fluctuating conditions). The comparative ease of gaining or losing copy number enables rapid toggling between ecological strategies (**Figure 5D**).

## Discussion

In this study, we extended our previously established CNV-reporter system in *S. cerevisiae* to simultaneously track copy number dynamics at two distinct genomic loci. Building upon existing strains containing the mCitrine gene adjacent to *GAP1*, we inserted the mCherry gene adjacent to additional nitrogen transporter genes known to undergo amplification under static nitrogen-limited conditions. Specifically, we generated CNV reporters for the proline transporter gene, *PUT4* and the ammonium-sulfate transporter gene, *MEP2*, both of which exhibit copy number variation in response to limitation for specific nitrogen source. This experimental design enabled single-cell analysis of copy number states at two chromosomal loci simultaneously, facilitating comprehensive and high-resolution analysis of population dynamics under both static and fluctuating environmental conditions.

Whereas previous research established a dual-fluorescent CNV reporter system for examining loss of heterozygosity at a single locus in the pathogenic diploid fungus *Candida albicans* (Zhou et al. 2024), our approach represents the first implementation of dual-fluorescent reporting for monitoring CNV dynamics at two distinct loci simultaneously. This methodological advancement enabled us to track multi-locus copy number variation allowing characterization of both genotype-by-environment (G × E) interactions and higher-order epistatic relationships (G × G × E).

Using this experimental system, we found that in static nitrogen-limiting environments evolutionary trajectories exhibited greater reproducibility across replicate populations compared to fluctuating conditions, especially in early stages. Consistent with previous investigations, *GAP1* amplifications demonstrated predictable selective sweeps through populations under glutamine-limitation, typically emerging around 70 generations (Lauer et al. 2018; Avecilla et al. 2022; Chuong et al. 2025). Whereas *PUT4* gene amplification has been documented during adaptive evolution in proline-limited environments, the temporal dynamics of *PUT4* amplifications had not been previously characterized (Hong and Gresham 2014). Although *PUT4* amplifications underwent positive selection in all replicate populations, two populations exhibited CNV frequency reversals, with *PUT4* copy number returning to single copy levels. This reversal coincided temporally with the emergence of potentially beneficial *GAP1* deletions. It is possible that *PUT4* amplifications may become subject to negative selection at later evolutionary stages due to diminishing returns associated with higher copy numbers. Whereas moderate amplifications (e.g., three copies) may confer a competitive advantage, increased amplification (e.g., five copies) potentially imposes metabolic burdens or regulatory costs that outweigh fitness benefits. Alternatively, *GAP1* deletions may provide superior fitness advantages compared to *PUT4* amplifications during later evolutionary phases. In this case, the fitness benefit of such deletions likely depends on the accumulation of background mutations that alter the fitness landscape explaining their absence during initial adaptation phases. These results are consistent with extensive experimental evidence demonstrating consistency in adaptive outcomes under stable conditions (Bennett, Lenski, and Mittler 1992; Bull et al. 1997; Cuevas, Elena, and Moya 2002; Tenaillon et al. 2012; Lauer et al. 2018; Chuong et al. 2025). Notably, the distinct evolutionary behaviors of these three nitrogen transporter genes demonstrate the locus-dependent nature of adaptive CNV dynamics underscoring the importance of multi-locus approaches for understanding adaptive evolution.

By contrast, fluctuating conditions fundamentally altered evolutionary dynamics, generating divergent outcomes between replicate populations. In environments alternating between glutamine- and proline-limitation, populations either showed no detectable *GAP1* or *PUT4* copy number changes, suggesting adaptation through alternative mechanisms of genetic variation, or adaptation through amplification of transporter genes at two loci simultaneously. This variability in outcome contrasts sharply with the reproducibility of static conditions and demonstrates that environmental fluctuations reduce evolutionary predictability (Cooper and Lenski 2010).

We propose that a specialist-generalist framework provides valuable insights into resource utilization strategies across diverse environments. While this concept is traditionally defined by phenotypic outcomes, increased fitness in single versus multiple environments, the underlying genetic variation remains poorly understood. Progress has been made in inferring ecological preferences from genomic data, such as the association between smaller genome sizes and specialized species versus larger genomes and generalists (Nancy A. Moran and Plague 2004; Klasson and Andersson 2004; N. A. Moran and Wernegreen 2000). However, these patterns primarily differentiate species rather than individuals within populations, highlighting the need for experimental models linking genomic variation to ecological function.

Previous studies of mutations in a specialization-generalization context have focused predominantly on SNVs, yet CNVs differ fundamentally in three critical aspects. First, CNVs arise through distinct mechanisms, including non-allelic homologous recombination (NAHR) (Peng et al. 2015), non-homologous end joining (NHEJ) (F. Zhang et al. 2009), and DNA replication errors (Brewer et al. 2011). Second, CNVs produce more predictable phenotypic outcomes compared to the context-dependent effects of SNVs. Third, CNVs exhibit continuous, proportional relationships between copy number and functional effect (Spealman et al. 2025), contrasting with the discrete effects of SNVs. These distinctive properties of CNVs may make them unique for inference of ecological strategies.

Our experimental results demonstrate that CNVs can mediate specialization and generalization strategies. Single-locus analysis revealed that increasing copy numbers enhanced specialization for both *GAP1* and *PUT4* independently. However, understanding generalization required simultaneous evaluation at two loci. Given the proportional relationship between gene copy number and functional effect, we hypothesized that generalists would exhibit equal copy numbers at both loci (0:0, 1:1, 2:2, 3:3). Fitness assessments across glutamine- and proline-limitation environments support this prediction. Genotypes with balanced copy numbers performed equally well in both environments, demonstrating generalist behavior, whereas specialists displayed asymmetric copy numbers with enhanced performance in the environment corresponding to the higher-copy locus. We find that static conditions favored selection for specialists, whereas fluctuating environments promoted selection for generalists. This trend is evident in both laboratory evolution experiments and in a survey of natural genetic variation. These findings underscore the unique capacity of CNVs to mediate rapid, reversible transitions between resource utilization strategies.

Taken together, our study shows that contrary to our initial expectation that CNVs would facilitate adaptive tracking, CNVs instead enable rapid, reversible shifts in ecological strategy, between generalist and specialist profiles, through changes in gene copy number balance across distinct resource-use axes. This supports a model in which CNVs do not function as genomic toggles between ecological states particularly in response to broad transitions between stable and fluctuating regimes. These findings suggest that genome architecture, specifically the modularity and reversibility of CNVs, can modulate the predictability of evolutionary outcomes by constraining or expanding access to distinct ecological niches.

Our study provides the first empirical demonstration that copy number ratios at nutrient transporter loci encode ecological strategy in microbial populations. By integrating experimental evolution with genomics data of thousands of natural isolates, we show that copy number symmetry marks generalists found in both wild and domesticated strains, whereas copy number asymmetry identifies specialists, which are restricted to stable, human-associated environments. This genomic signature of niche breadth offers a new framework for linking structural variation to ecological function. The principles established here may extend broadly to other microbes, where CNVs at transporter loci underlie ecological plasticity. By framing CNV balance as a molecular determinant of ecological strategy, this work opens new avenues for predicting functional traits from genome structure, informing the study of resource-use tradeoffs, life history evolution, and adaptive responses in dynamic environments.

## Methods

### Media and growth conditions

Nitrogen-limiting media (glutamine, proline, and ammonium-sulfate) contained 800 μM nitrogen regardless of molecular form and 1 g/L CaCl_2_-2H_2_O, 1 g/L of NaCl, 5 g/L of MgSO_4_-7H_2_O, 10 g/L KH_2_PO_4_, 2% glucose, and trace metals and vitamins as previously described (Hong and Gresham 2014).

### Strain construction

We used haploid FY4 derivatives of the reference strain S288c, for all experiments. To generate dual-fluorescent CNV-reporter strains, we used the CNV-reporter strain, DGY1584, generated in (Lauer et al. 2018) with the constitutively expressed *mCitrine* gene marked by the kanMX G418-resistance cassette (*TEFpr*::*kanMX*::*TEFterm*) inserted adjacent to the *GAP1* locus as our base strain.

Using Gibson assembly, we constructed the plasmid DGP363, encoding *prACT1*-*mCherry*-*HygR*-*ADH1term* We PCR-amplified the mCherry cassette using primers with 40bp homology and performed high-efficiency yeast transformation (Gietz and Schiestl 2007) to insert the construct adjacent to either *PUT4* (DGY2318;integration coordinate: Chromosome XV: 989384), or *MEP2* (DGY2323;integration coordinate: Chromosome XIV: 359195). For 1-copy control strains, DGY500, DGY2347, and DGY2349, either the mCitrine or mCherry reporter was integrated at one of two neutral loci: either replacing *HO* (*YDL227C*) on Chromosome IV or replacing the dubious ORFs, *YLR122C* and *YLR123C*, on Chromosome XII. To generate either 2-copy mCitrine or mCherry haploid controls, we mated the 1-copy controls, sporulated and dissected the resulting diploids to identify segregants containing both CNV reporters (DGY1315 for mCitrine and DGY2331 for mCherry). PCR and Sanger sequencing confirmed correct integration of all CNV reporters(strains and primer sequences used in this study are provided in Supplementary table 1).

### Long-term experimental evolution

Starter cultures were prepared by streaking engineered strains from frozen glycerol stocks onto YPD plates and incubating them overnight at 30°C. A single colony from each plate was used to inoculate batch cultures in nitrogen-limiting medium, each with the relevant nitrogen source, and then grown overnight at 30°C. Once the cultures had reached sufficient density, cell counts were measured with a Coulter Counter to ensure comparable starting densities. We transferred 0.3 mL of each dual-fluorescent CNV reporter culture into 20-mL ministat vessels (Miller et al. 2013) and allowed them to grow overnight in batch mode in a 30°C growth chamber, to reach the desired steady-state cell density. After confirming that all cultures had reached similar cell densities, the pumps were activated, switching the system from batch mode to continuous culture. For static conditions either glutamine-, ammonium sulfate-, or proline-limited media was continuously pumped into vessels from 10 L carboys. For fluctuating conditions, we manually switched media type by connecting two media carboys to each chemostat vessel and using plastic clamps to close one inlet and open the other. Control populations containing either one or two copies of the CNV reporter replacing neutral loci (*HO* and *YLR122/23C*) were also inoculated in ministat vessels for each media condition. Ministats were maintained at 30°C in aerobic conditions and diluted at a rate of 0.12 hour^−1^ (corresponding to a population doubling time of 5.8 hours). Steady-state populations of 3 × 10^7^ cells were maintained in continuous mode for 250 generations (60 days). Every 10-20 generations, with the exception of a larger gap between generation 80 and 164 generations, we collected 1mL for flow cytometry analysis.

### Flow cytometry analysis of dual-CNV reporter system

To track CNV dynamics at the *GAP1*, *PUT4*, and *MEP2* loci, we briefly sonicated samples to disrupt clumps and immediately analyzed them on a Cytek Aurora flow cytometer. For each sample, we recorded data from approximately 10,000 cells, capturing forward scatter (FSC), side scatter (SSC), and fluorescence intensity in both the mCitrine and mCherry channels. For comparing mean fluorescence between mCitrine and mCherry, we normalized the fluorescence values for the two fluorophores using the three reference thresholds of the 0-, 1-, 2-copies of mCitrine and mCherry to determine the appropriate scaling factor. For determining copy number per single-cell, we employed hierarchical gating to define key subpopulations: cells, singlets, and cells with varying mCitrine and mCherry copy numbers. First, we gated for cells by plotting FSC-A vs SSC-A to exclude debris and microbial contamination. To evaluate the impact of static versus fluctuating conditions on cell physiology, we compared cell size between these environments and found that, in most cases, there were no significant differences (**Supplementary figure 2**). Next, we gated for single cells by plotting FSC-A vs FSC-H and selecting the diagonal population to exclude doublets. Finally, we plotted mCitrine fluorescence (excitation 488 nm, emission 530 nm) against mCherry fluorescence (excitation 561 nm, emission 610 nm) to distinguish copy number states. Gates were manually drawn to define populations with zero, one, or multiple copies of each fluorophore, using zero-, single-, and two-copy controls as references. Cells with three or more copies were defined as all events with fluorescence intensity exceeding the upper boundary of the two-copy gate. While we observed partially separated clusters at higher fluorescence levels, potentially corresponding to four or five gene copies, we conservatively grouped these cells into a single three-plus category due to the lack of validated controls for higher copy number states.

### Muller plot generation

Using the single-cell genotype frequencies inferred from flow cytometry, we constructed Muller plots to visualize the dynamics of CNV genotypes over time. For each population and locus, cells were assigned to discrete genotypic classes (zero, one, two, and three-plus copies), and their population frequency at each time point was calculated. To account for minor overlap between adjacent fluorescence gates, we applied a correction in which 10% of the measured 2-copy population was subtracted and reallocated to the 1-copy class at each time point.

Negative values resulting from this adjustment were clipped to zero, and the deficit was added back to the 1-copy class to preserve total abundance. Muller plots were generated using the R package ggmuller (Noble and Golding 2017), with lineage relationships assigned manually. The ancestral 1-copy genotype was designated as the parent, and higher or lower copy number classes were treated as descendants. These visualizations captured CNV-driven clonal expansions and genotype turnover across environments.

### Modelling genotype-by-environment and epistatic interactions

Fitness values were obtained by first binning counts for each pairwise combination of *GAP1* copy number and either *PUT4* or *MEP2* copy number, ranging from zero to three. Log fold change was then calculated between generation four, the earliest time point sampled, and generation 260, the final time point sampled, as follows:

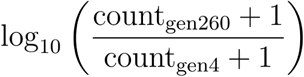

A pseudocount of one was added to cell counts to account for samples with zero recorded events, ensuring defined values across all comparisons and avoiding infinite values from division by zero. This approach resulted in four distinct values corresponding to each replicate and denoting 0 - 3+ copies of the secondary locus, decreasing the likelihood that our model would be susceptible to random noise from a small number of values. To ensure that the bins used had sufficient observations we assessed the number of cells in each bin across experiments (**Supplementary figure 9**).

To assess whether the effect of mCherry copy number on fitness varied across different mCitrine copy number states, we performed a linear regression analysis within each medium condition. Specifically, we modeled the log-transformed fold change in cell counts as a function of an interaction between mCherry and mCitrine copy number. mCitrine copy number was treated as a categorical variable with four levels (0, 1, 2, and 3), using copy number 1 as the reference level. The interaction terms between mCherry copy number (treated as a continuous variable) and each non-reference mCitrine copy number level were extracted and interpreted as the differential effect of mCherry copy number on fitness in those mCitrine backgrounds. Confidence intervals for model coefficients were calculated to evaluate statistical support.

### Fitness calculations for determining fitness profiles

To quantify fitness effects associated with different copy number ratios in glutamine-limitation and proline-limitation, we binned data by mCitrine and mCherry copy number ratio and summed counts over all replicates in a particular condition. This allowed us first to average over variations that may have been attributed to low counts for some CNV ratio groups and second to have respective values for each CNV ratio group in both glutamine- and proline-limitation.

Fitness was calculated as the log-transformed fold change in cell number across time:

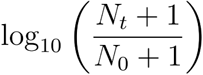

where *N*_0_ and *N*□ represent estimated cell numbers at generations 4 and 260, respectively. A pseudocount of 1 was added to avoid undefined values for genotypes with zero abundance at either generation.

### Determining gene copy number and ecological strategy of natural strains

We used CNV estimates generated by CNVnator from the 3,034 *S. cerevisiae* genomes published by Loegler et al. (2024). Briefly, they calculated median normalized read depth for each gene locus genome-wide. Loci were assigned the computed copy number if a CNV was detected, otherwise, a default value of 1 was assigned. To focus specifically on variation in copy number, we excluded all strains with a value of 1 at a given locus, as these were not expected to contribute to CNV-driven differences. To convert continuous normalized read depth values into discrete copy number estimates, we rounded each value up to the nearest integer.

Ecological strategy was then assigned based on the ratio of copy numbers at two gene loci: strains with highly unequal copy numbers were classified as specialists, while those with similar copy numbers at both loci were classified as generalists.

## Acknowledgements

We thank previous and present members of the Gresham, Vogel and Ghedin labs and Conor Gilligan for valuable discussion and comments on the manuscript. We thank the NYU Genomics Core facility and specifically Hana Husic for flow cytometry services. We acknowledge the Zegar Family Foundation for their generous support. Funding for this research was provided by NIGMS (R35GM153419) and the US - Israel Binational Science Foundation (2021276).

---

**Supplementary figure 1.**
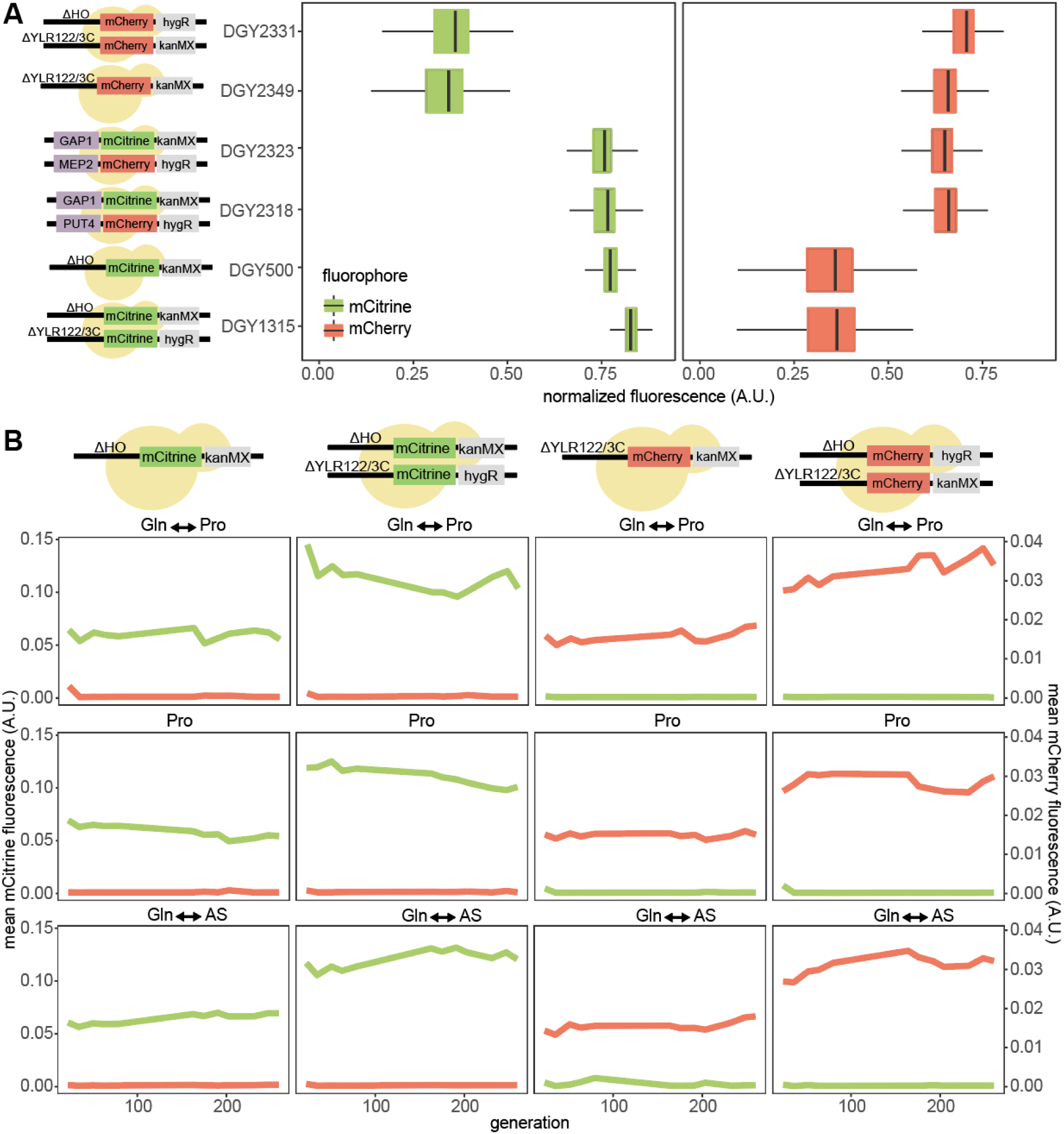
Establishing a dual-fluorescence CNV reporter. Boxplots showing the normalized fluorescence profiles of the constructed dual-fluorescent strains for both mCitrine and mCherry including both control and experimental strains (A). Control strains experimentally evolved in three distinct conditions for 260 generations. Control strains include either 1 copy mCitrine, 1 copy mCherry, 2 copies mCitrine, or 2 copies of mCherry. The red line represents mCherry fluorescence levels, and the green line represents mCitrine fluorescence levels (B).

**Supplementary figure 2.**
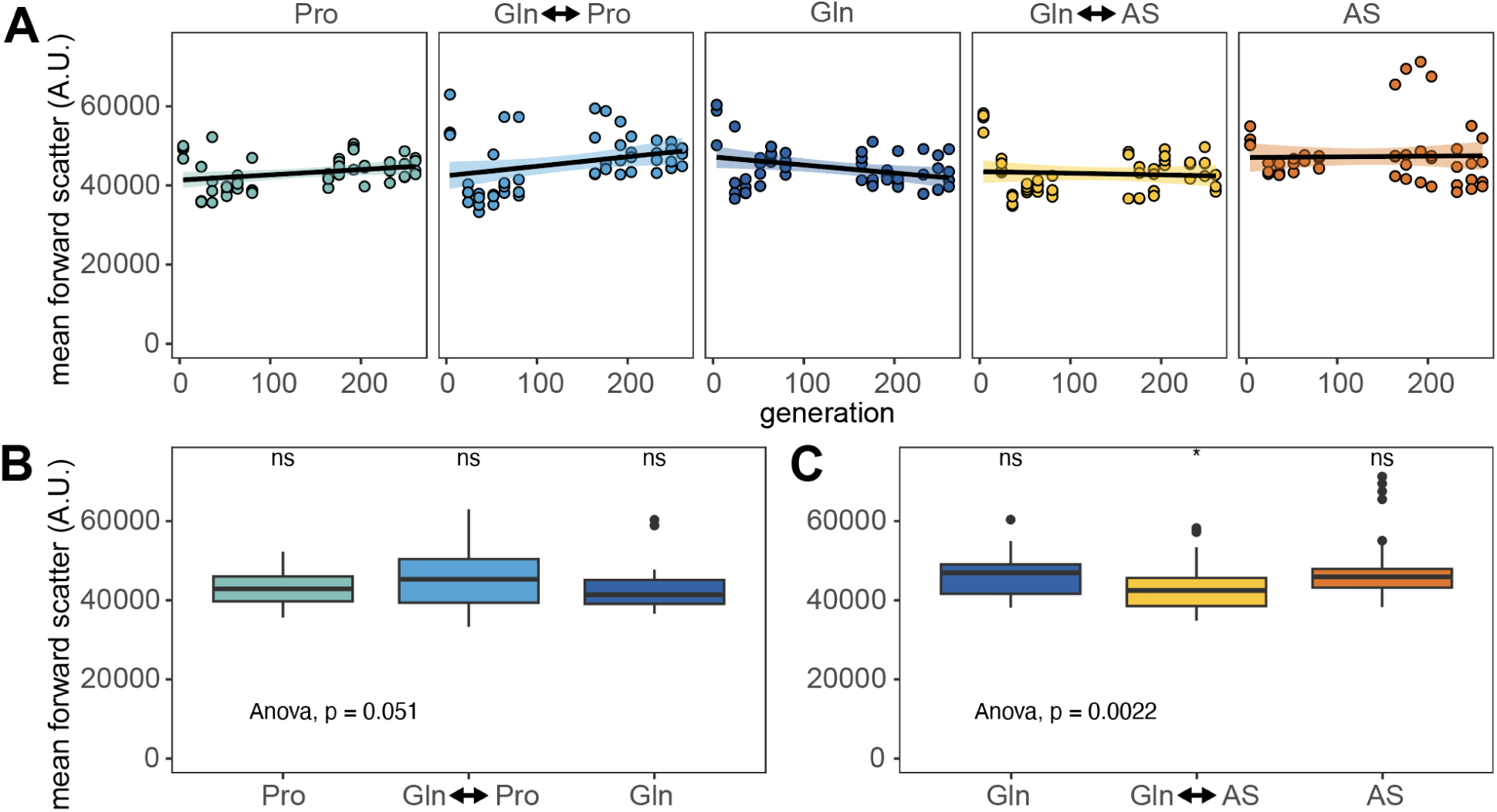
Forward scatter, a proxy measurement for cell size, generally shows little significant change across conditions. (A) Mean forward scatter across experimental evolution time is shown for each condition which is made up for replicate populations. (B) Boxplots of mean forward scatter for static proline-limitation (Pro) and glutamine-limitation (Gln) and fluctuating between proline-limitation and glutamine-limitation with the anova test performed pairwise for all three conditions and a t-test to determine significance. (C) Boxplots of mean forward scatter for static ammonium-sulfate-limitation (AS) and glutamine-limitation (Gln) and fluctuating between proline-limitation and glutamine-limitation with the anova test performed pairwise for all three conditions and a t-test to determine significance.

**Supplementary figure 3.**
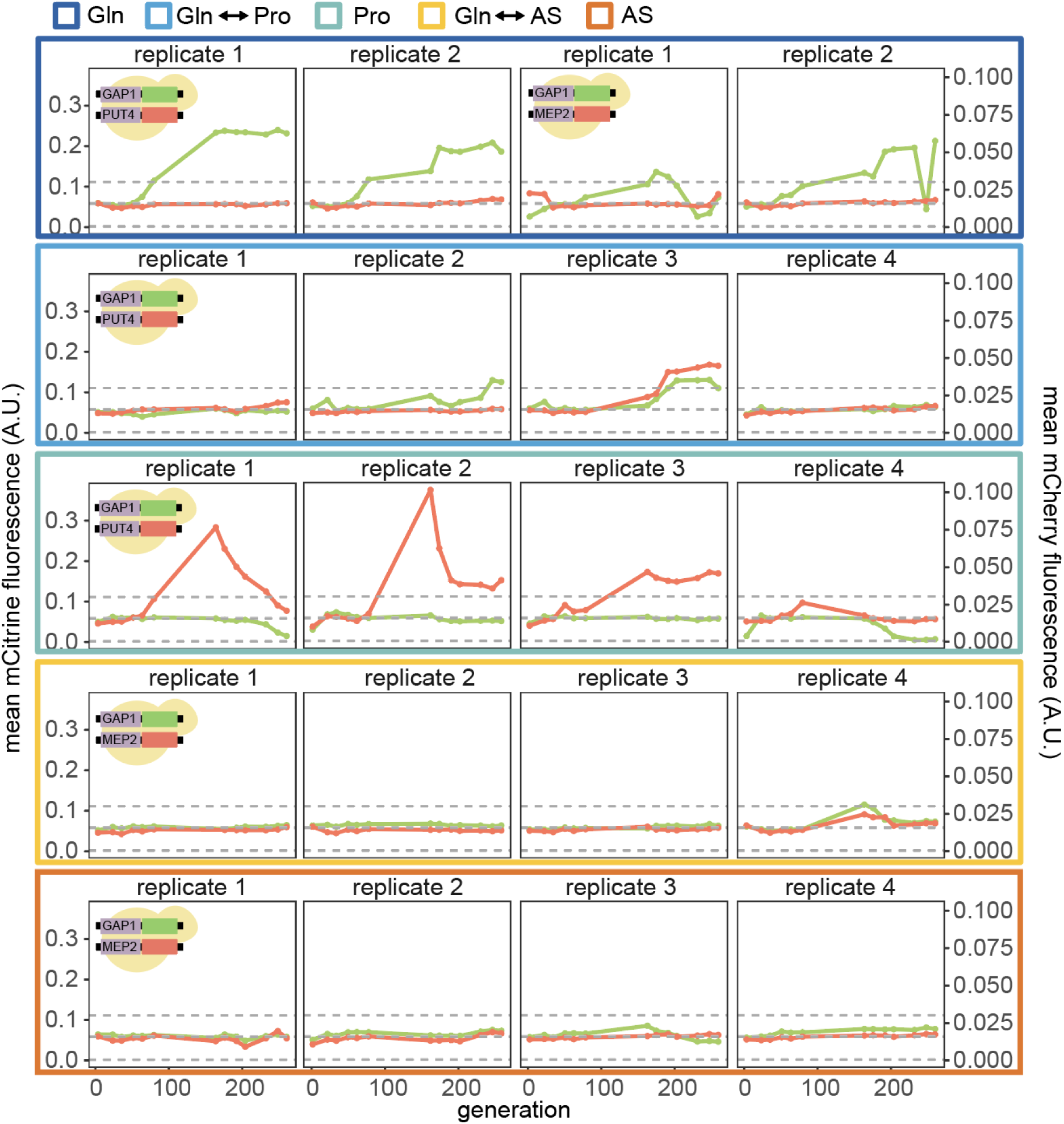
Population dynamics of copy number variation across conditions. Mean fluorescence profiles for both mCitrine and mCherry dynamics across all five conditions. The three gray dashed lines represent the mean fluorescence value of the 0-, 1-, and 2- copy control strains from bottom to top.

**Supplementary figure 4.**
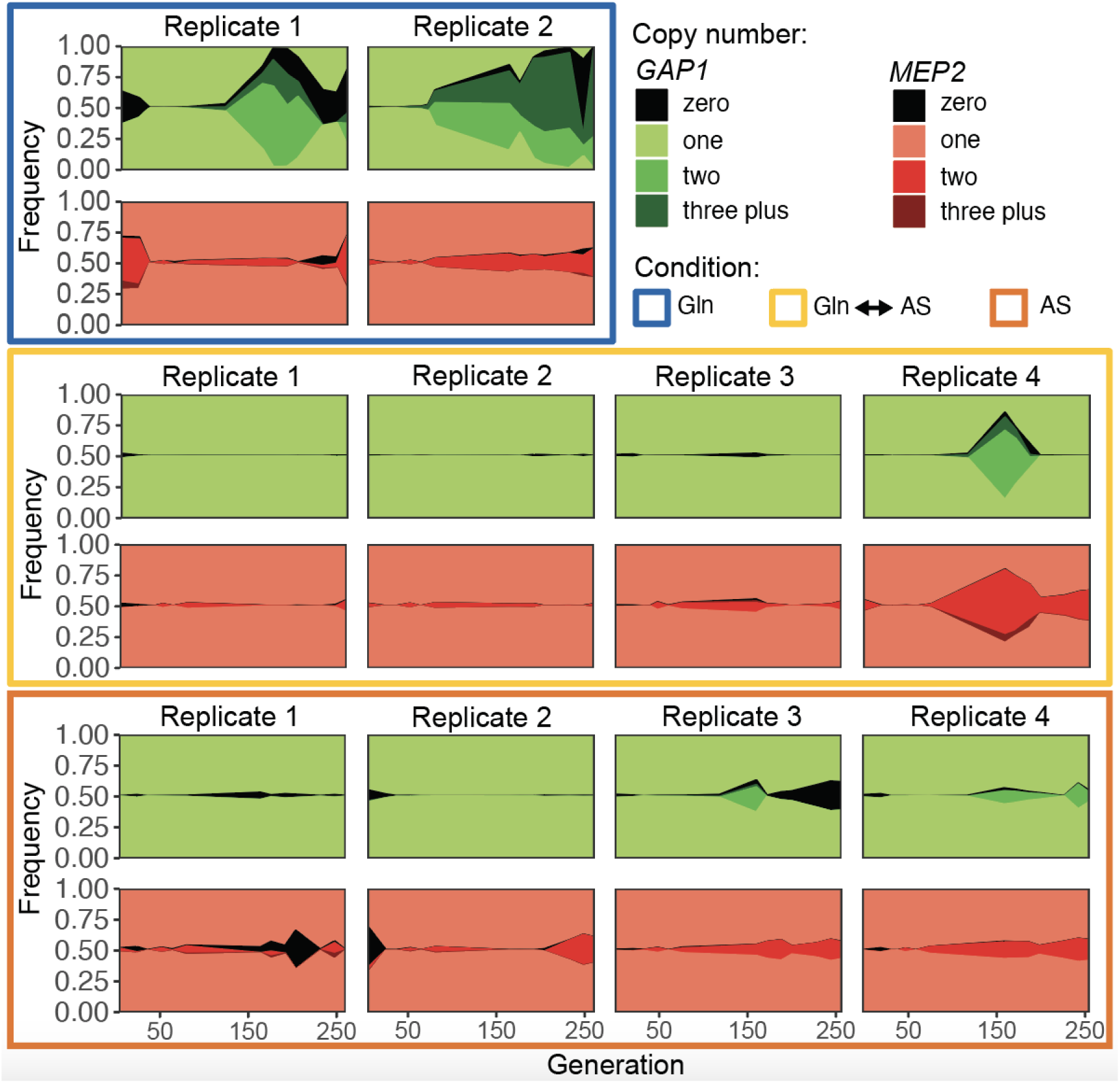
Single-cell CNV dynamics of the *GAP1*/*MEP2* CNV reporter strain across conditions. Muller plots are used to visualize the proportion of cells containing zero, one, two and three plus copy number of either *GAP1* or *MEP2* the static conditions glutamine-limitation and ammonium sulfate-limitation and the condition that fluctuates between the two.

**Supplementary figure 5.**
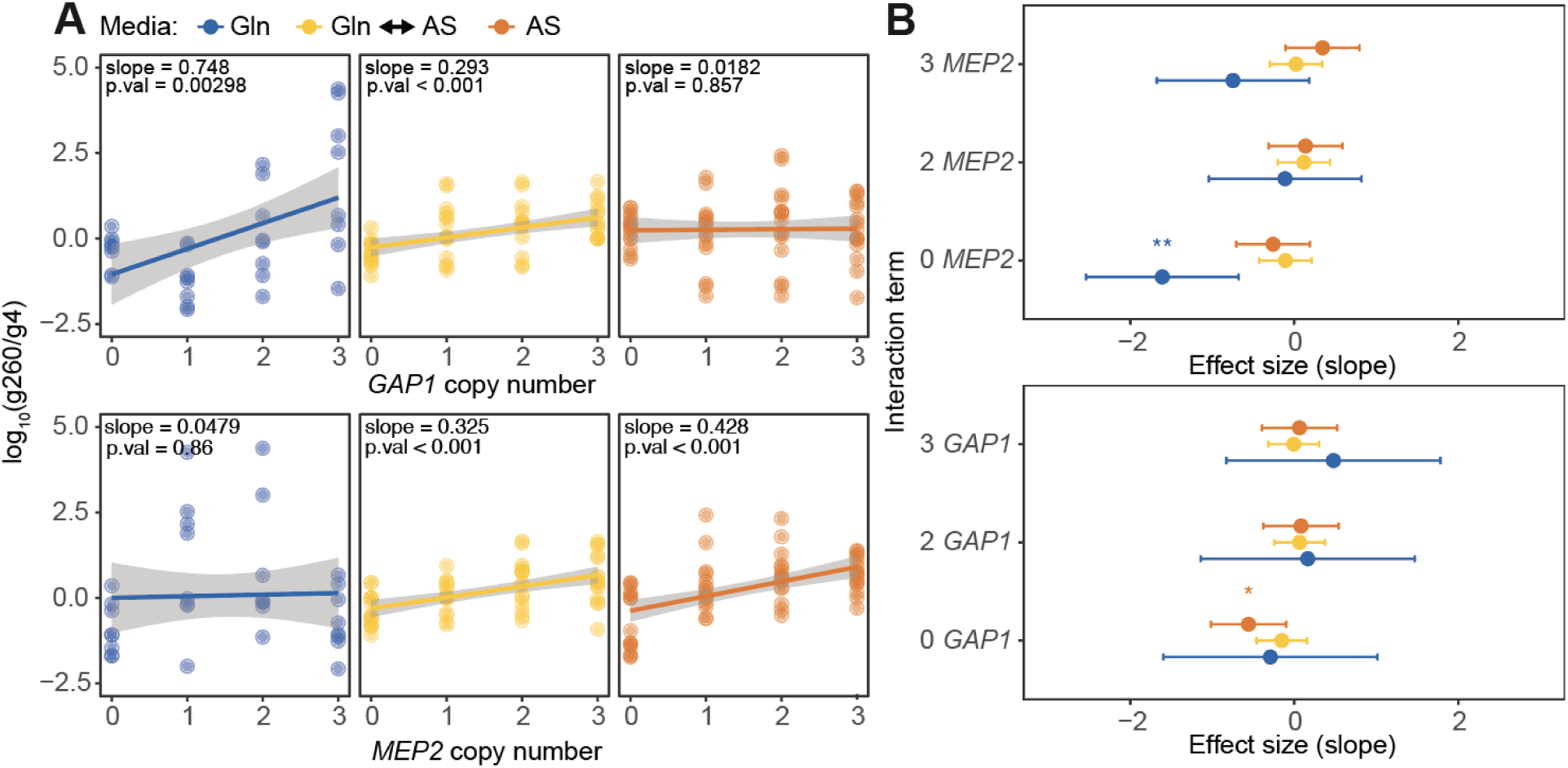
Quantifying environmental and genetic interactions of *GAP1* and *MEP2* using the *GAP1/MEP2* CNV reporter strain. Testing how fitness scales with *GAP1* (top) and *MEP2* (bottom) copy number in different environments (G × E) (A). Testing whether copy number values of the secondary locus interact with the primary locus to alter fitness (G × G × E). P-value was annotated with an asterisk as follows: ‘*’ for <0.05, ‘**’ for < 0.01, ‘***’ for < 0.001 (B).

**Supplementary figure 6.**
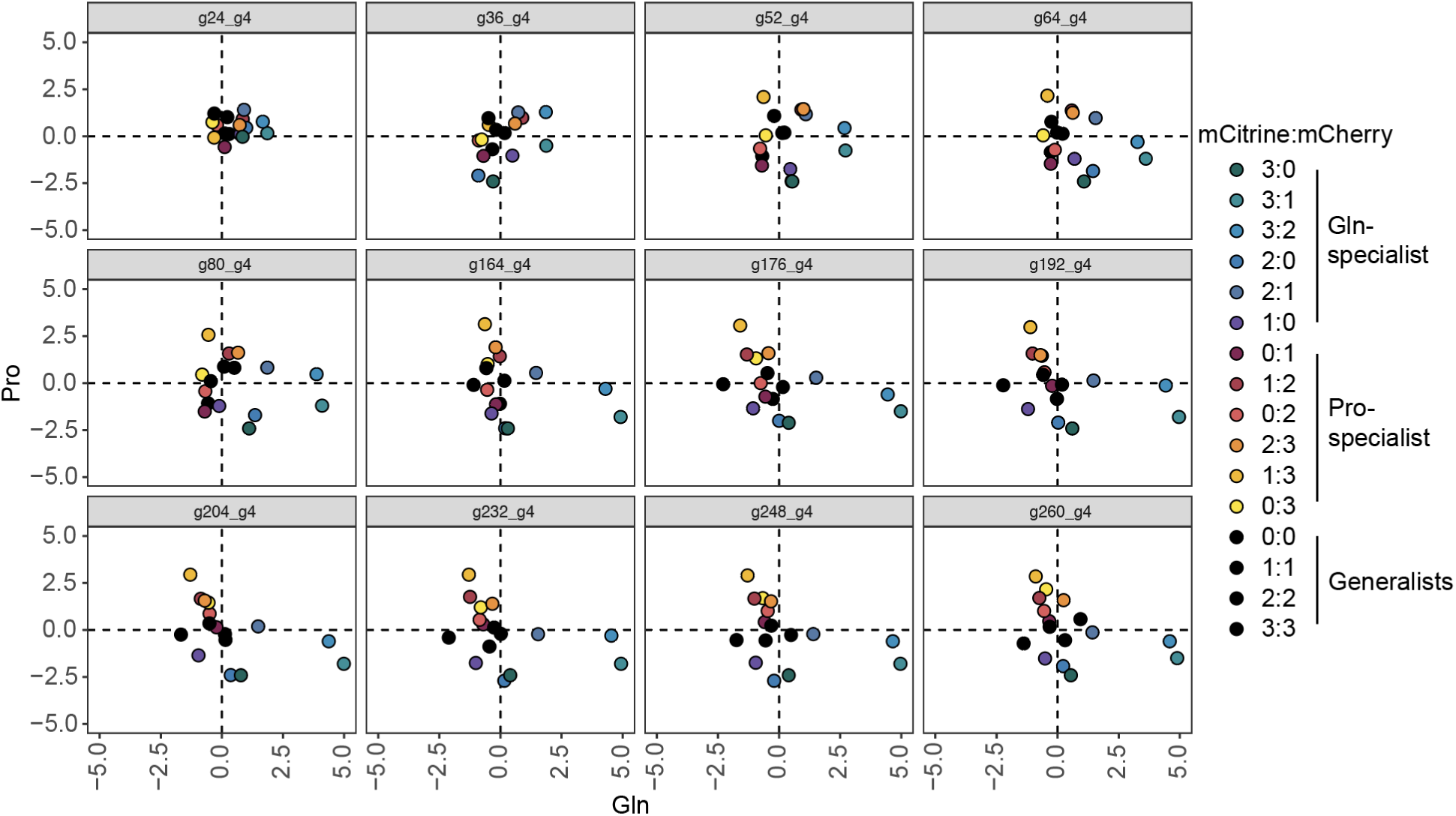
Relationship between fitness under proline vs. glutamine-limitation, with fitness being calculated across all generations relative to generation 4. Points are colored based on the *GAP1/PUT4* ratio.

**Supplementary figure 7.**
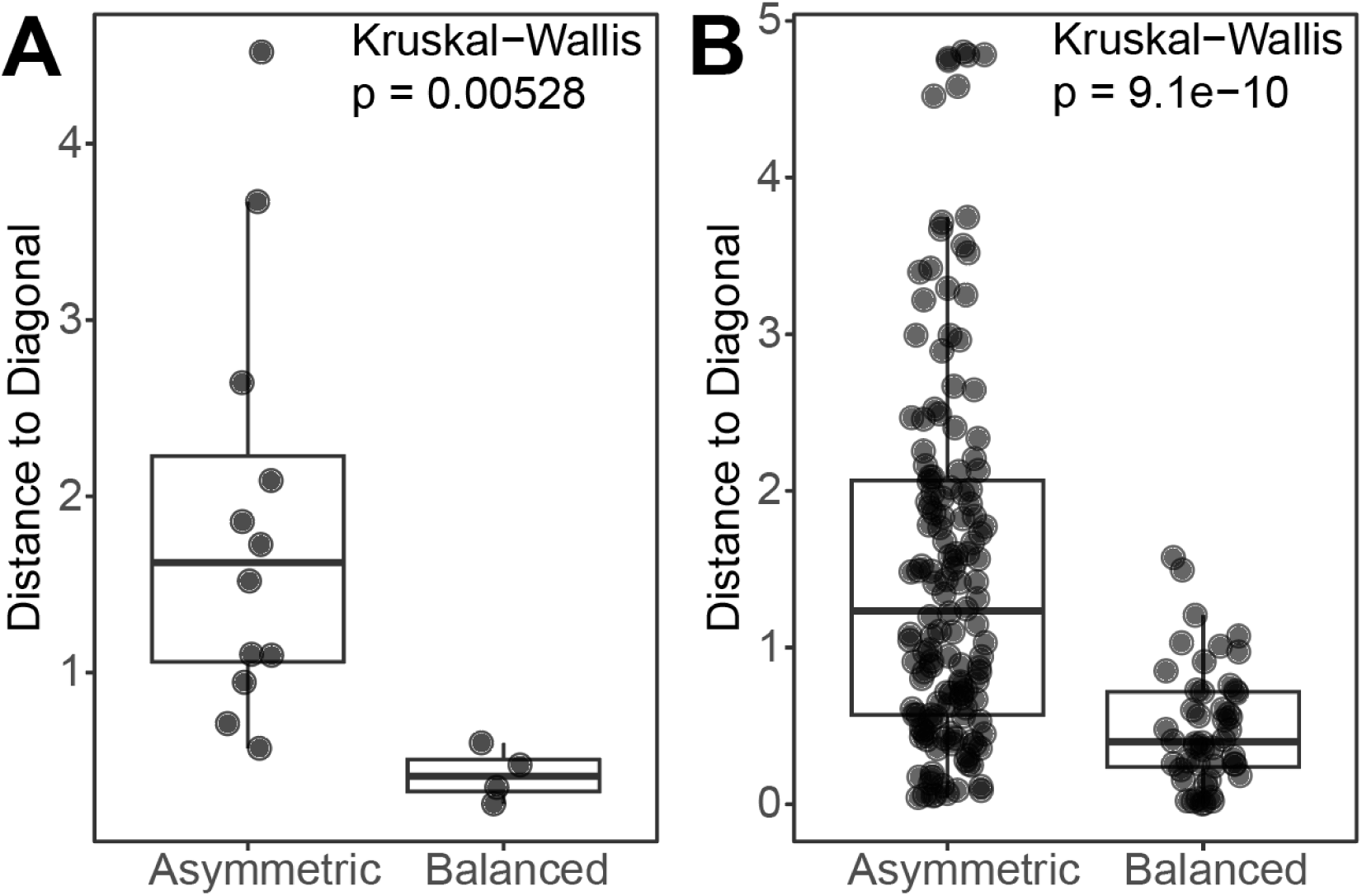
Balanced copy number genotypes cluster near the diagonal (x = y), consistent with expectations for generalists. Distance to the diagonal was calculated as the 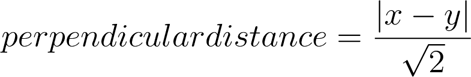. Fitness was computed as the log_10_ ratio of counts between generation 260 and generation 4 (A). Perpendicular distance for fitness values across time was calculated from all pairwise log_10_ ratios between each generation and generation 4 (B). Statistical significance was assessed using the Kruskal–Wallis test (p = 0.05).

**Supplementary figure 8.**
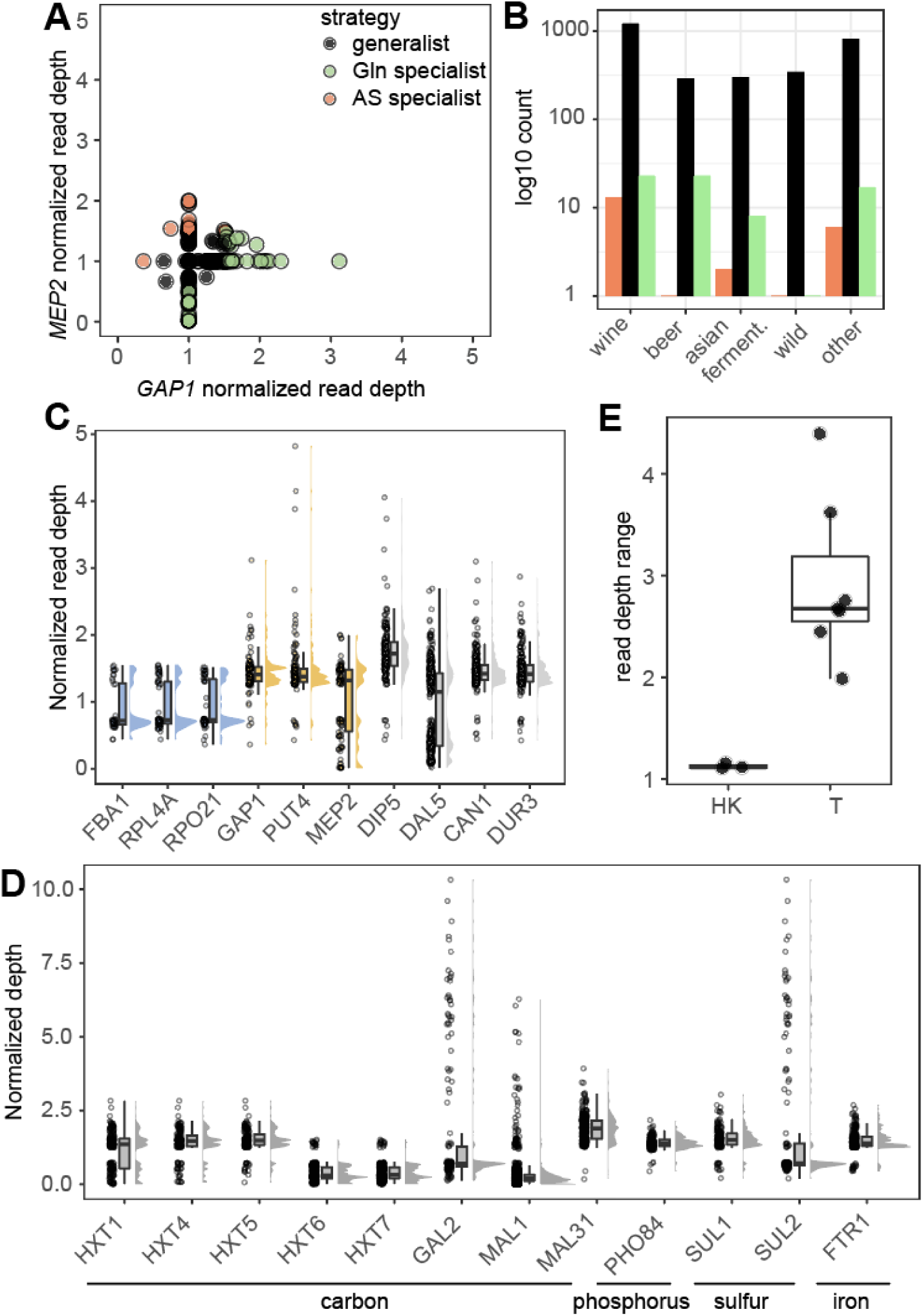
Various transporters show greater copy number variation than housekeeping genes in natural isolates. Normalized read depth for *MEP2* and *GAP1* as a proxy for gene copy number with ecological strategy assignments (A). Barplots showing the number of strains falling within each ecological strategy across yeast isolate super clades (B). Normalized read depth of housekeeping genes in blue, nitrogen transporter genes assessed by the dual-fluorescence system are in gold and additional transporter genes not assessed in this manuscript are in gray (C). Normalized read depth of transporters of additional substrates (D). Boxplots comparing read depth range of housekeeping genes (HK) to transporter genes (T) shown in plots C and D (E).

**Supplementary figure 9.**
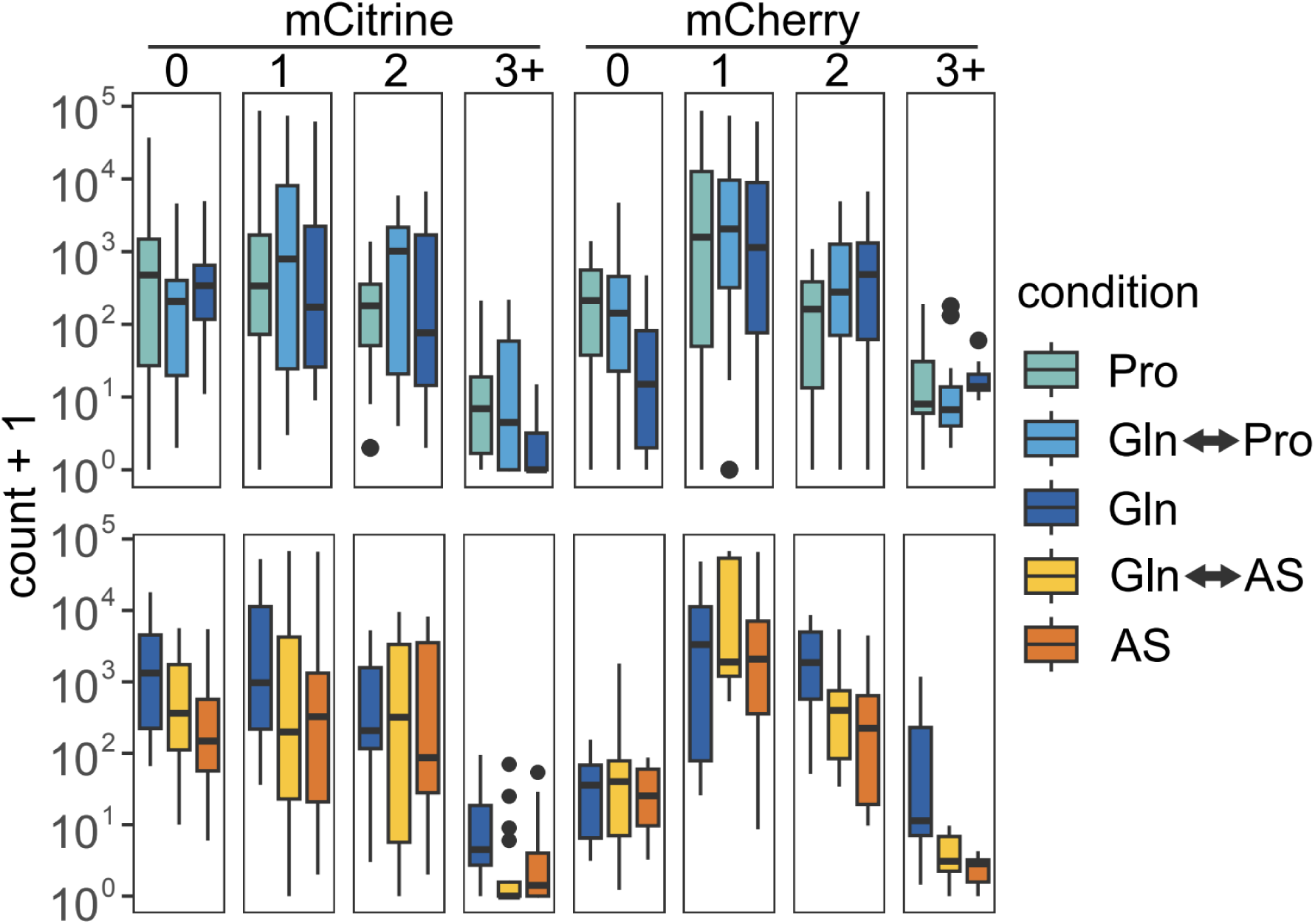
Boxplots of cell counts of each copy number bin for mCitrine and mCherry at the first sampled timepoint, generation four. As expected, the greatest cell count is of the one copy bin since it is the ancestral genotype. The three plus copy bin has the lowest counts. Samples plotted or static and fluctuating conditions for either the *GAP1/PUT4* reporter strain (top) or the *GAP1/MEP2* reporter strain (bottom).

**Supplementary table 1.**
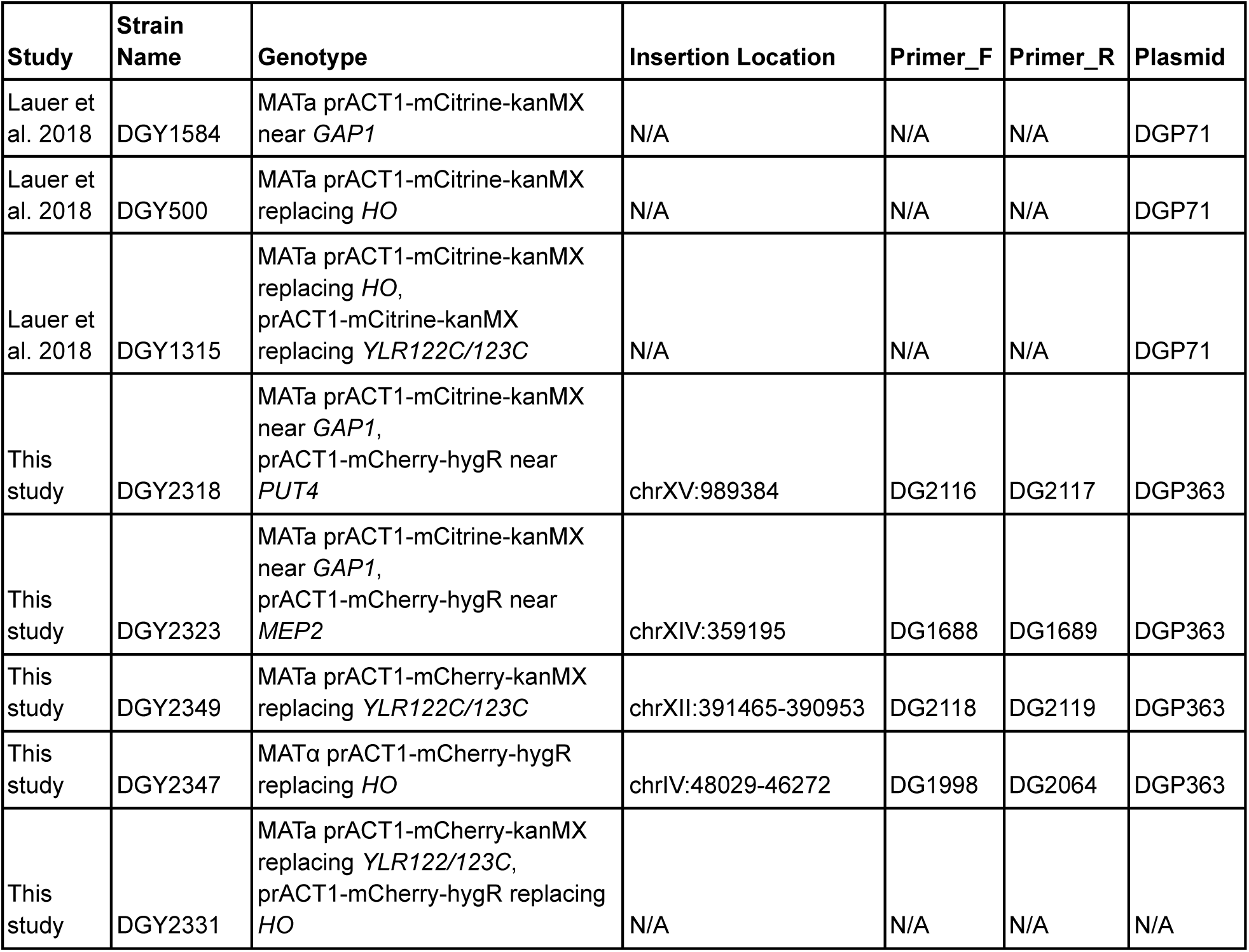
*S. cerevisiae* strains used in this study. Strains were either previously described in Lauer et al. (2018) or newly constructed for this study. For newly constructed strains, the corresponding plasmid and primers used are listed, along with the chromosomal integration site of the inserted cassette.

